# *Zbtb46* coordinates angiogenesis and immunity to control tumor outcome

**DOI:** 10.1101/2023.03.02.530835

**Authors:** Ashraf Ul Kabir, Madhav Subramanian, Jun Wu, Minseo Kim, Karen Krchma, Xiaoli Wang, Carmen M. Halabi, Hua Pan, Samuel A. Wickline, Daved H. Fremont, Kyunghee Choi

## Abstract

Tumor-angiogenesis and -immunity play critical roles in cancer progression and outcome. An inverse correlation of these two^1^ hints at common regulatory mechanism(s). Here, we report that *Zbtb46*, a repressive transcription factor and a widely accepted marker for classical dendritic cells (DCs)^2, 3^, constitutes one such regulatory mechanism. *Zbtb46* was downregulated in both DCs and endothelial cells (ECs) by tumor-derived factors to facilitate robust tumor growth. *Zbtb46* downregulation led to a hallmark pro-tumor microenvironment (TME), including dysfunctional vasculature and immunosuppressive cell accumulation. Analysis of cancer patient data revealed a similar association of low *ZBTB46* expression with an immunosuppressive TME and a worse prognosis. In contrast, enforced *Zbtb46* expression mitigated the pro-tumor TME features and restricted tumor growth. Mechanistically, *Zbtb46-*deficient ECs were highly angiogenic, and *Zbtb46-*deficient bone-marrow progenitors upregulated *Cebpb* and diverted the DC program to myeloid lineage output, potentially explaining the myeloid lineage skewing phenomenon in cancer^4–7^. Conversely, enforced *Zbtb46* expression normalized tumor vessels and, by suppressing *Cebpb*, skewed bone-marrow precursors towards more DC generation over macrophages, leading to an immune-hot TME. Remarkably, *Zbtb46* mRNA treatment synergized with anti-PD1 immunotherapy to improve tumor management in pre-clinical models. These findings identify *Zbtb46* as a common regulatory mechanism for angiogenesis and for myeloid lineage skewing in cancer and suggest that maintaining its expression could have therapeutic benefits.

## Main

The cooperation between tumor cells and the tumor microenvironment (TME) is a defining characteristic of cancer^8, 9^. As such, targeting the communication between the tumor and TME has recently gained much attention^10, 11^. One such strategy is immune checkpoint blockade (ICB) immunotherapy, which has significantly impacted the cancer treatment landscape by promoting long-term tumor control and complete regression in some patients^12–14^. Unfortunately, most patients still do not respond to ICB therapies^15, 16^. The efficacy of ICB relies on the ability of T cells to kill cancer cells and the support of the vasculature and antigen-presenting cells, such as dendritic cells (DCs), in delivering and activating these T cells, respectively. Tumors can disrupt this process by suppressing anti-tumor programs in the TME components^10^. Identifying these suppressive programs has tremendous potential for use as adjuvants to improve the outcomes of immunotherapies.

*Zbtb46* is a member of the BTB-ZF family of transcriptional repressors and is considered a marker of classical DCs^2, 3^. Endothelial cells (ECs) of the vascular system also constitutively express *Zbtb46*^2^. In homeostatic conditions, *Zbtb46* is thought to keep both these cell types in a quiescent state^17, 18^; however, its function has not been addressed in any pathological conditions. Here, we report that *Zbtb46* expression orchestrates a critical tumor-suppressor program in the vasculature and hematopoietic system. Tumor-derived factors downregulate *Zbtb46* expression, leading to a pro-tumor TME. Our study finds that maintaining *Zbtb46* expression in a therapeutic manner results in an anti-tumor TME and enhances the effectiveness of ICB treatment.

### *Zbtb46* is a negative regulator of tumor progression

ECs and classical DCs express *Zbtb46* constitutively in physiological conditions; however, *Zbtb46* deficiency in the healthy state does not lead to any discernible pathological conditions^2, 3^. We, too, observed that *Zbtb46^gfp/gfp^* (*Zbtb46* KO or ZKO) mice did not display any pathological complications related to the cardiovascular system in a homeostatic state (Extended Data Fig. 1a,b). We investigated the direct involvement of *Zbtb46* in solid tumor progression using mouse cancer models. *Zbtb46* expression was downregulated in the tumor-associated ECs and DCs in the wild-type (WT) mice bearing 1956 sarcoma^19, 20^, LLC carcinoma^21^, and PyMT-BO1^22^ (orthotopic) and MMTV-PyMT^23^ (genetic) breast cancers (Fig. 1a and Extended Data Fig. 1c-g). This suppression was also observed in the bone marrow (BM) cells (Fig. 1a). We challenged ZKO mice with the same tumor models to determine whether this suppression contributes to tumor outcome. Compared to their WT counterparts, ZKO mice exhibited more robust tumor growth (Fig. 1b-e). Further analysis revealed enhanced pro-tumor TME features in the ZKO mice, including exuberant angiogenesis, lower CTL/Treg and M1/M2 ratios, decreased cDC1 presence, and increased fibroblast activation (Fig. 1f,g and Extended Data Fig. 2a). Remarkably, analysis of the cancer patient survival data showed that a lower expression of *ZBTB46* is associated with a worse prognosis in multiple cancer types (Extended Data Fig. 2b). Similar to the animal models, patients with a lower expression of *ZBTB46* had reduced anti-tumor immune components (Fig. 1h and Extended Data Fig. 2c). Together, these data suggest a constitutive inhibitory role for *Zbtb46* in tumor growth.

**Fig.1:**
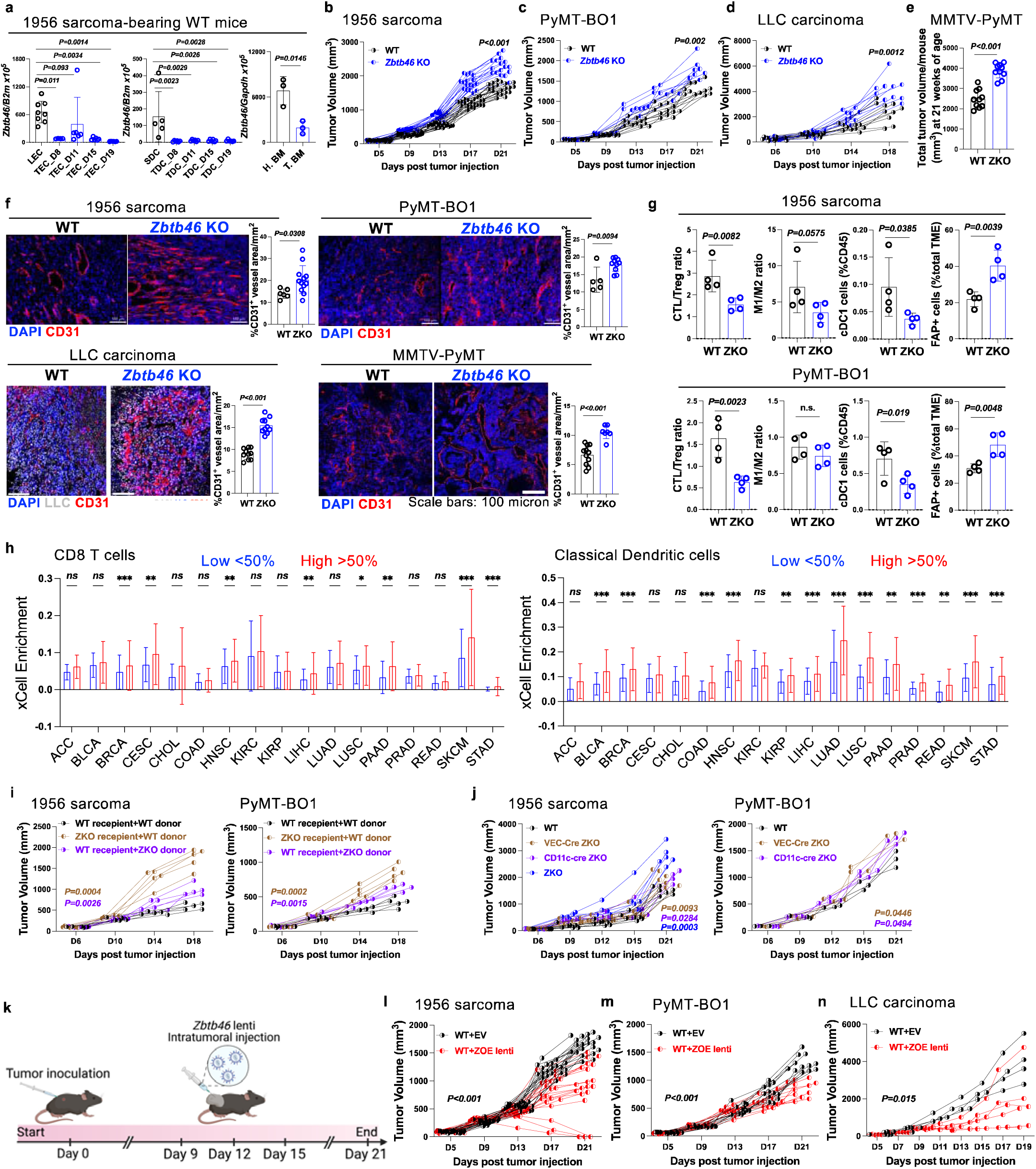
Zbtb46 is a negative regulator of tumor progression. **a**, *Zbtb46* mRNA expression in (left) CD31+CD45-ECs, (middle) CD45+CD11c+MHCII+ DCs, and (right) bone marrow cells isolated from the lungs (LEC), Spleen (SDC), and tibia and femur (H. BM) of the healthy wild-type mice and from the tumors (TEC and TDC) and tibia and femur (T. BM) of the 1956 sarcoma-bearing wild-type mice at indicated days post-transplantation. n≥3/group. Data are mean±SD. (left and middle) One-way ANOVA with Dunnett’s test and (right) Student’s t-test. **b-e**, Tumor growth in (b) 1956 sarcoma, (c) PyMT-BO1 breast cancer, (d) LLC carcinoma, and (e) MMTV-PyMT breast cancer in wild-type and *Zbtb46* KO mice. n≥8/group. Student’s t-test at endpoint. **f**, Representative images with quantification of tumoral CD31+ vessel density in wild-type and *Zbtb46* KO mice bearing indicated tumors. n≥4/group. Data are mean±SD. Student’s t-test. **g**, Measurements of CTL/Treg, M1/M2, CD11c+MHCII+XCR1+ cDC1 cells, and Fibroblast activation protein (FAP)+ in the TME of 1956 sarcoma and PyMT-BO1 breast tumor-bearing mice at endpoints described in b and c. n=4/group. Data are mean±SD. Student’s t-test. **h**, Presence of CD8T and classical dendritic cells in high vs. low *ZBTB46* expressing tumors in patients from the TCGA database analyzed with XCell. Data are mean±SEM. Student’s t-test. *p<0.05, **p<0.01, ***p<0.001, ns=not significant. **i**, Tumor growth of 1956 sarcoma and PyMT-BO1 breast cancer in stromal *Zbtb46* KO and hematopoietic *Zbtb46* KO mice. n≥5/group. One-way ANOVA with Dunnett’s test at endpoint compared to control mice. **j**, Tumor growth of 1956 sarcoma and PyMT-BO1 breast cancer in wild-type, VEC-cre *Zbtb46* KO, CD11c-cre *Zbtb46* KO, and *Zbtb46* KO mice. n≥4/group. One-way ANOVA with Dunnett’s test at endpoint compared to wild-type mice. **k-n**, (k) Schematics and tumor growth of (l) 1956 sarcoma, (m) PyMT-BO1 breast cancer, and (n) LLC carcinoma in wild-type mice with either empty vector (EV) or *Zbtb46* (ZOE) lentiviral overexpression construct (intra-tumor) treatment. n≥8/group. Student’s t-test at endpoint compared to EV-treated mice.

We examined the relative contribution of EC- and DC-specific *Zbtb46* expression in the tumor growth by employing endothelial (VEC-Cre)- or hematopoietic (CD11c-Cre or VAV-Cre)-specific deletion of *Zbtb46.* We also generated BM chimeras for the same purpose (Extended Data Fig. 3a). In both systems, we observed that *Zbtb46* was required in both the endothelial and hematopoietic cells for optimal tumor control (Fig. 1i,j and Extended Data Fig. 3b). The most robust tumor growth was observed in the global ZKO mice, suggesting a potential collaborative role for the EC- and DC-specific *Zbtb46* expression in suppressing tumor growth.

If *Zbtb46* acted as a tumor-suppressor in the TME that tumors target to downregulate, an enforced expression could halt tumor progression. Indeed, intra-tumor lentiviral delivery of *Zbtb46* led to sustained expression in the TME and reduced tumor mass (Fig. 1k-n and Extended Data Fig. 3c). Importantly, *Zbtb46* overexpressing tumor cell lines did not have much growth difference compared to the parental tumor cell lines, suggesting a TME-centric role for *Zbtb46* expression in suppressing tumor growth (Extended Data Fig. 3d).

### *Zbtb46* normalizes tumor vasculature

Tumor vessels are inherently abnormal, featuring impaired vascular perfusion and excessive leakage that leads to a hypoxic and immunosuppressive microenvironment^24, 25^. Along with the tumor growth restriction, intra-tumor lentiviral *Zbtb46* expression curbed tumor angiogenesis (Fig. 2a). While tumor vasculature was more dysfunctional in the ZKO mice, intra-tumor enforced *Zbtb46* expression improved all the functional characteristics leading to vascular normalization (Fig. 2b and Extended Data Fig. 4a; also see scheme in Fig. 1k). Normalized tumor vessels are known to support more anti-tumor immune and stromal TME^26–30^, which could partly explain the decreased anti-tumor immune components observed in the ZKO mice and the opposite following the enforced *Zbtb46* expression (Fig. 1g; also see Fig. 3c). Similar to a previous report^18^, we found that *Zbtb46* overexpression in mouse cardiac endothelial cells (MCEC) made them quiescent, as evidenced by modestly reduced proliferation and significantly limited tube-formation capabilities in vitro (Extended Data Fig. 4b,c). Bulk mRNA sequencing revealed that while the parental MCEC cells were enriched in migration and angiogenic pathways, *Zbtb46* overexpressing MCEC cells were enriched in hypoxic stress response and immune-supportive pathways (Fig. 2c). Moreover, *Zbtb46* overexpressing MCEC cells allowed more leukocyte trans-endothelial migration, another hallmark feature of the normalized vasculature^29^ (Fig. 2d). Finally, we established a co-transplantation system consisting of ECs and tumor cells, where the co-transplanted ECs directly impact tumor growth. Remarkably, EC-specific *Zbtb46* overexpression significantly restricted the tumor progression, solidifying the importance of *Zbtb46* expression in the tumor vasculature (Fig. 2e).

**Fig.2:**
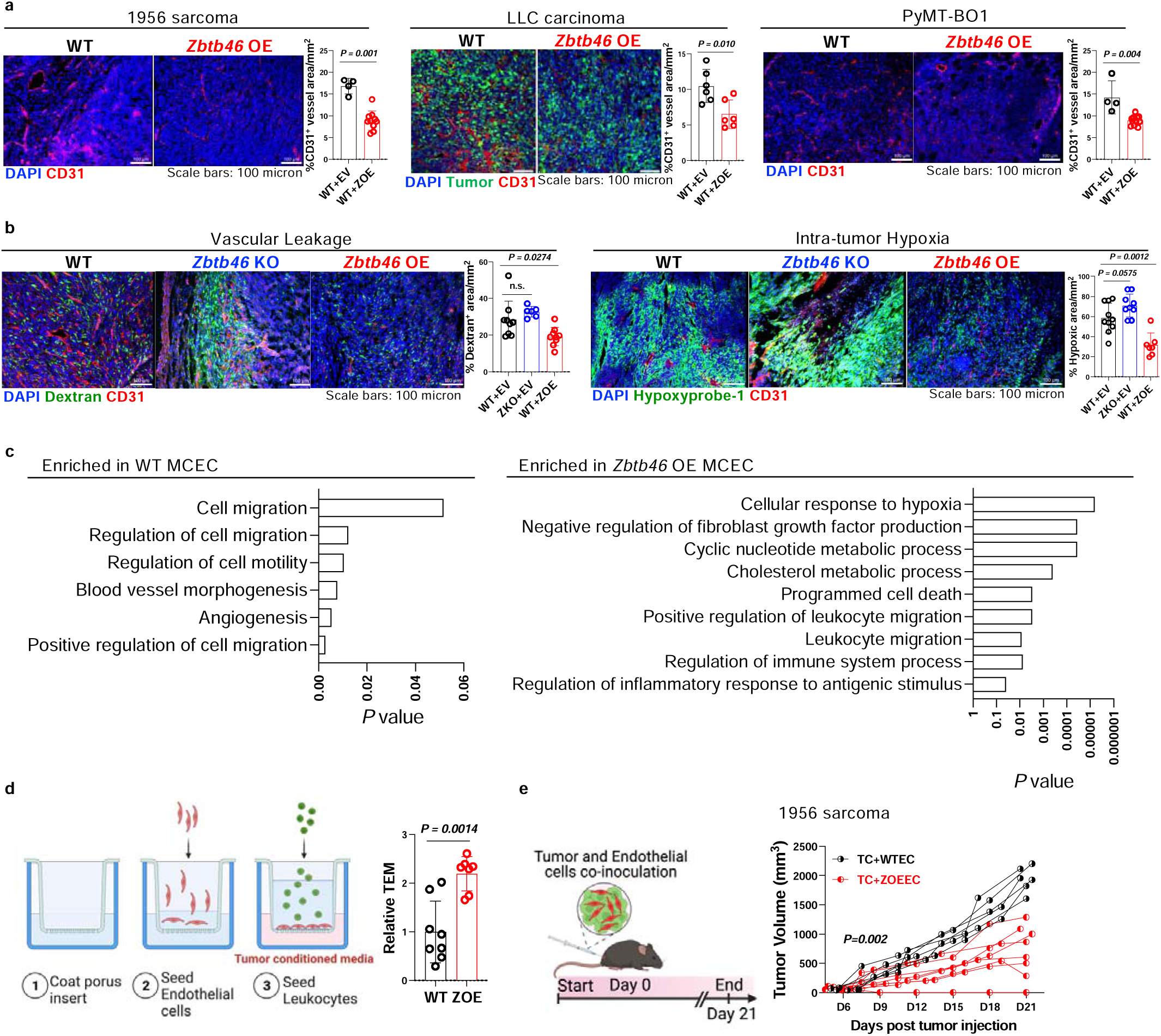
Zbtb46 normalizes tumor vasculature. **a**, Representative images with quantification of tumoral CD31+ vessel density in wild-type mice with either empty vector (WT+EV) or *Zbtb46* (WT+ZOE) lentiviral overexpression construct (intra-tumor) treatment bearing indicated tumors. n≥4/group. Data are mean±SD. Student’s t-test. **b**, Representative images and quantification for vascular leakage and intra-tumoral hypoxia as measured by the FITC-Dextran 70KD spread and the relative abundance of Hypoxyprobe-1 binding, respectively, in the 1956 sarcoma tumor tissue of wild-type mice with either empty vector (WT+EV) or *Zbtb46* (WT+ZOE) or of *Zbtb46* KO mice with empty vector (ZKO+EV) lentiviral overexpression construct (intra-tumor) treatment. n≥4/group. Data are mean±SD. One-way ANOVA with Dunnett’s test. **c**, GO enrichment analysis of bulk-RNA sequencing data from parental (WT) and *Zbtb46*-overexpressed (OE) MCEC cells. **d**, Schematic and analysis of relative migration of CD3+CD8+ CTL cells through parental (WT) and *Zbtb46* overexpressed (ZOE) MCEC cell barrier in a leukocyte trans-endothelial migration assay. Data are mean±SD. Student’s t-test. **e**, Schematics and tumor growth of 1956 sarcoma cells co-transplanted with either parental ECs (TC+WTEC) or *Zbtb46*-overexpressed ECs (TC+ZOEEC). n≥6/group. Student’s t-test at endpoint.

**Fig.3:**
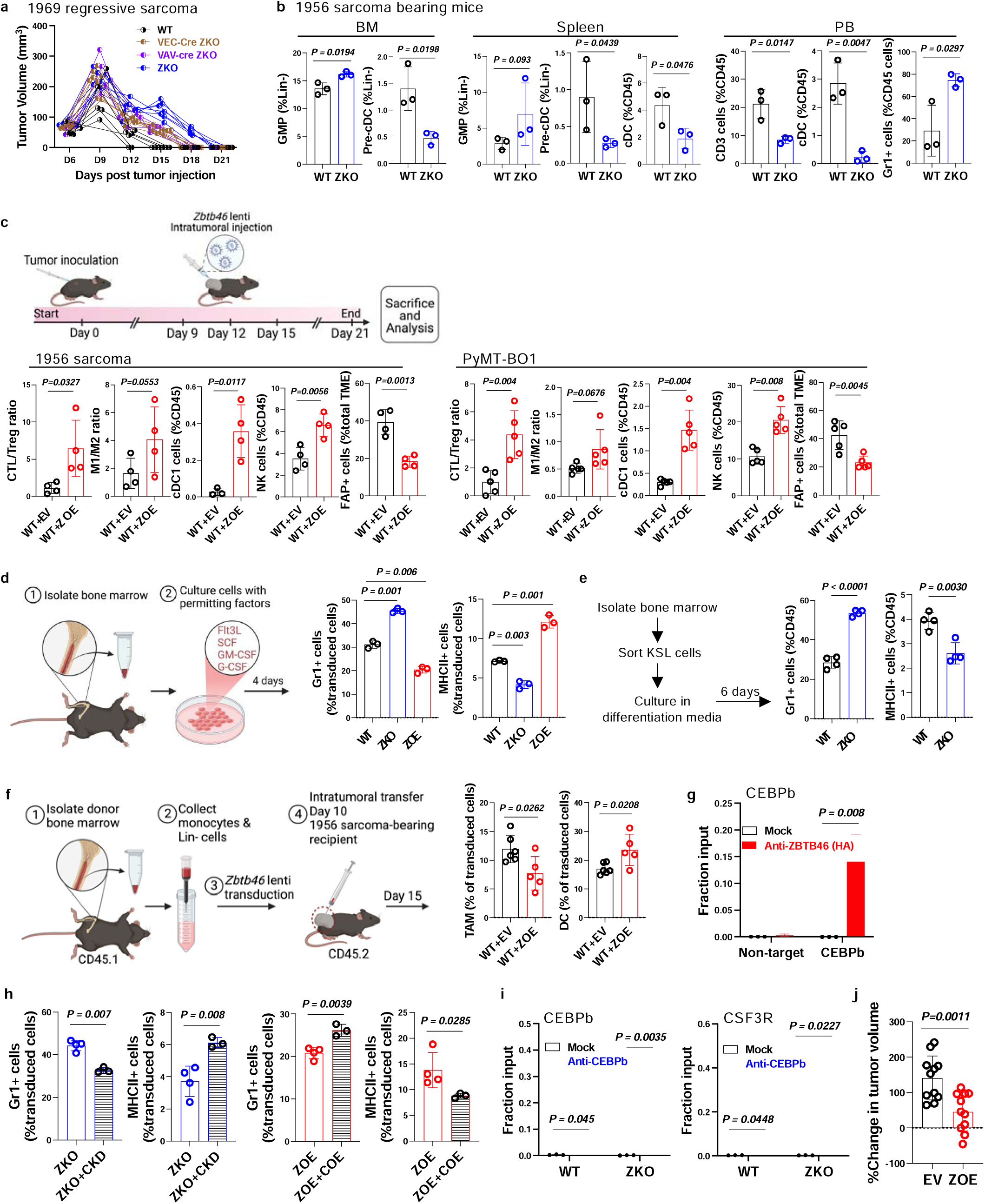
Zbtb46 promotes an anti-tumor hematopoietic system. **a**, Tumor growth of 1969 regressive sarcoma in wild-type, VEC-cre *Zbtb46* KO, VAV-cre *Zbtb46* KO, and *Zbtb46* KO mice. n≥6/group. **b**, Analysis of GMP, Pre-cDC, cDC, CD3, and Gr1+ cells in the bone marrow (BM), spleen, and peripheral blood (PB) of the 1956 sarcoma-bearing mice as indicated. GMP: Lin-Sca1-cKit+CD41-CD150-CD16/32+, Pre-cDC: Lin-Sca1-IL7R-MHCII-CD16/32-CD11c+cKit-CD135+, cDC: CD11c+MHCII+. n≥3/group. Data are mean±SD. Student’s t-test. **c**, Analysis of the tumor immune microenvironment of 1956 sarcoma-bearing and PyMT-BO1 tumor-bearing wild-type mice with either empty vector (EV) or *Zbtb46* (ZOE) lentiviral overexpression construct (intra-tumor) treatment. n≥4/group. Data are mean±SD. Student’s t-test. **d**, Analysis of Gr1+ and MHCII+ cell generation from wild-type or *Zbtb46* KO mice bone marrow cells with either empty vector-mCherry (WT and ZKO) or *Zbtb46*-mCherry (ZOE) lentiviral overexpression. n=3/group. Data are mean±SD. One-way ANOVA with Dunnett’s test. **e**, Analysis of Gr1+ and MHCII+ cell generation from wild-type or *Zbtb46* KO mice KSL cells (Lin-Sca1+cKit+) sorted from bone marrow. n=4/group. Data are mean±SD. Student’s t-test. **f**, Analysis of donor-derived tumor-associated macrophages (TAM) and DCs after five days of intra-tumor transfer of enriched monocytes from wild-type donor mice bone marrow (CD45.1) into 1956 sarcoma-bearing wild-type recipient mice (CD45.2) with either empty vector-mCherry (WT+EV) or *Zbtb46*-mCherry (WT+ZOE) lentiviral overexpression. n≥5/group. Data are mean±SD. Student’s t-test. **g**, ChIP-qPCR analysis of ZBTB46 recruitment to a potential binding site in *Cebpb* promoter region. *Zbtb46*-HA lentiviral construct was overexpressed in wild-type mice bone marrow-derived precursors (monocyte and lineage-negative cells) and ChIP was performed with an anti-HA antibody. n≥3/group. Data are mean±SD. Student’s t-test. **h**, Analysis of Gr1+ and MHCII+ cell generation from bone marrow cells of *Zbtb46* KO mice with empty vector-mCherry lentiviral overexpression alone (ZKO) or with *Cebpb* shRNA constructs (ZKO+CKD) and of wild-type mice with *Zbtb46*-mCherry lentiviral overexpression alone (ZOE) or with *Cebpb* overexpression (ZOE+COE). n=3/group. This experiment is a part of the experiment described in **d**. Data are mean±SD. Student’s t-test. **i**, ChIP-qPCR analysis for CEBPB enrichment at CEBPB peaks in bone marrow-derived precursors (monocyte and lineage-negative cells) from wild-type or *Zbtb46* KO mice. n≥3/group. Data are mean±SD. Student’s t-test. **j**, Analysis of tumor progression after five days of intra-tumoral transfer of enriched monocytes from wild-type mice bone marrow into 1956 sarcoma-bearing wild-type mice with either empty vector (EV) or *Zbtb46* (ZOE) lentiviral overexpression. n≥6/group. Data are mean±SD. Student’s t-test.

### *Zbtb46* promotes DC generation while restricting myeloid lineage output

*Zbtb46* deficiency does not compromise classical DC function in a steady state^2, 3, 17^. To assess whether the scenario is different in cancer, we challenged the global, EC-specific (VEC-Cre), and hematopoietic-specific (CD11c-Cre or VAV-Cre) ZKO mice with 1969 sarcoma^19, 20^ tumors that regress spontaneously in immune-competent WT mice. Strikingly, all the ZKO mice rejected the inoculated 1969 tumor mass, albeit with a slightly delayed kinetics compared to the WT mice (Fig. 3a and Extended Data Fig. 5a). This outcome is unlike what was reported for *Batf3* KO mice, a gene critical for cDC1 lineage development and cDC1-mediated cross-presentation for tumor rejection^31, 32^, suggesting an absence of any direct involvement of *Zbtb46* in classical DC function.

While *Zbtb46* KO classical DCs do not show any functional defects under tumor challenge, our data suggested that *Zbtb46* expression is required in the hematopoietic compartment for optimal tumor control (see Fig. 1i,j). We determined if there might be any other immune system-related changes by the *Zbtb46* expression status. Like the vascular system, the hematopoietic system in ZKO mice was mostly similar to the WT mice in homeostatic condition when we investigated both the progenitor- and committed-cell types in lymphoid organs and peripheral blood, except with a slight increase in the Gr1^+^ cells in the PB (Extended Data Fig. 5b). However, upon tumor challenge, the hematopoietic system in ZKO mice acquired more pronounced myeloid-biased pro-tumor immune characteristics as evidenced by the increased granulocyte-monocyte progenitor (GMP) and reduced pre-cDCs in both BM and spleen, as well as decreased T cells and cDCs and increased Gr1^+^ cells in the systemic circulation (Fig. 3b). In contrast, intra-tumor lentiviral *Zbtb46* delivery, which restricted tumor growth and angiogenesis, led to a more anti-tumor immune microenvironment, reflected by the increased CTL/Treg and M1/M2 ratios, enhanced cDC1 and NK cell population, and reduced fibroblast activation in the TME (Fig. 3c).

To further understand the impact of *Zbtb46* on the immune outcome, we isolated BM cells and cultured them with a mixture of factors supporting DC and myeloid lineage development^2^. *Zbtb46* deficient BM cells generated reduced MHCII^+^ DCs and increased Gr1^+^ myeloid cells compared to the WT BM cells (Fig. 3d). ZKO KSL (cKit^+^Sca1^+^Lin^-^) cell population that contains hematopoietic stem and progenitor cells (HSPC) produced more Gr1^+^ cells and fewer MHCII^+^ DCs compared to the WT KSL cells, suggesting that the myeloid lineage skewing occurs at a more primitive progenitor level (Fig. 3e). Conversely, enforced *Zbtb46* expression in BM cells reversed this trend to generate more DCs and fewer myeloid cells (Fig. 3d), in line with a previous observation^2^. More importantly, enforced *Zbtb46* expression in BM-derived progenitors, mostly containing monocytes and lineage-negative cells (see methods), led to the generation of more DCs and fewer macrophages when transferred intra-tumor into an established tumor, a system that closely reflects the tumor-infiltrating monocyte differentiation^33^ (Fig. 3f).

Our findings suggested that *Zbtb46* expression can control DC vs. myeloid differentiation output. Careful examination of the previous ZBTB46 ChIP-Seq^17^ and microarray^2^ data identified critical myeloid genes, including *Cebpb*, a crucial transcription factor for emergency granulopoiesis^34^ and monocyte/macrophage gene regulation^35^, as potential direct targets of *ZBTB46*. By utilizing ChIP-qPCR, we confirmed that ZBTB46 could bind the *Cebpb* promoter region in BM-derived progenitors (Fig. 3g and Extended Data Fig. 6a). While *Cebpb* expression was upregulated in the ZKO BM cells, enforced *Zbtb46* expression downregulated *Cebpb* (Extended Data Fig. 6b). Moreover, overexpression or knockdown of *Cebpb* partially reversed the overexpression or knockdown effect of *Zbtb46*, respectively, on myeloid and DC lineage generation from BM cells (Fig. 3h and Extended Data Fig. 6c). These observations and the fact that the consensus DNA recognition site for both ZBTB46 and CEBPB is highly similar^17, 36^ (Extended Data Fig. 6d) raise the possibility that *Cebpb* can further upregulate myeloid genes that are targets of ZBTB46 repression in the absence of *Zbtb46*. Indeed, in the same cell system from ZKO mice, CEBPB binding to its’ transcriptional targets, such as *Cebpb* itself and *Csf3r*, is significantly enhanced compared to the cells from WT mice (Fig. 3i and Extended Data Fig. 6e). A reporter system consisting of *Cebpb* transcriptional response element (TRE) (containing tandem repeats of consensus DNA recognition motifs) further demonstrated that *Zbtb46* can reduce the CEBPB-induced activation of TRE (Extended Data Fig. 6f,g). This interplay between *Zbtb46* and *Cebpb* is functionally important as BM cells from tumor-challenged ZKO mice had higher macrophage gene expression and lower DC gene expression than the WT tumor-challenged mice; enforced *Zbtb46* expression reversed this expression pattern (Extended Data Fig. 6h,i). These data suggest that the *Zbtb46* regulation of *Cebpb* is at least partly responsible for the enforced *Zbtb46*-mediated immunostimulatory TME. Similar to the EC-specific overexpression outcome, intra-tumor transfer of *Zbtb46*-overexpressed BM-derived progenitors had partially restricted tumor growth (Fig. 3j), strengthening the notion of the collaborative nature of the cell-type-specific *Zbtb46* expression in the tumor.

### Therapeutic maintenance of *Zbtb46* improves cancer immunotherapy outcome

Our data so far suggested that enforced *Zbtb46* expression in the TME led to vascular normalization and more DC generation with enhanced anti-tumor immunity. Because an immune-hot TME is a prerequisite for effective ICB therapy^1, 12^, we assessed whether the *Zbtb46*-maintenance could synergize with anti-PD1 immunotherapy. First, we generated *Zbtb46* mRNA nanoparticle using the p5RHH peptide system. The p5RHH peptide-based nanoparticle system readily formulates mRNA for systemic administration and extrahepatic nucleotide delivery and showed promise for safe and effective clinical translation^21, 29, 37–41^. Systemic administration of *Zbtb46* mRNA nanoparticle was effective in sustaining *Zbtb46* expression in tumor-DCs and -ECs and resulted in the restriction of tumor growth (Extended Data Fig. 7a-g). Although tumor cells also acquired *Zbtb46* expression (Extended Data Fig. 7c), as shown above, tumor cells overexpressing *Zbtb46* do not show much growth differences compared to parental lines (see Extended Data Fig. 3d), suggesting the EC- and DC-focused *Zbtb46* delivery as the driver for the observed tumor control. The tumor growth restriction was associated with an immunostimulatory TME (Extended Data Fig. 7a-g). Remarkably, the nanoparticles induced dramatic response in both anti-PD1-responsive (1956 sarcoma) and -refractory (PyMT-BO1) tumor^29^ models following the treatment, generating long-term complete remission of tumor mass in many of the treated animals challenged with the anti-PD1-responsive 1956 sarcoma tumor (Fig. 4a-d). The addition of a VEGFR2 inhibitor (DC101) enhanced the treatment response hinting at the collaborative nature of the targeted *Zbtb46* and *Vegf* pathways in the tumor (Fig. 4a-d). The tumor-eliminated mice spontaneously rejected the secondary challenge with the same tumor, indicative of the development of a long-term immunological memory (Extended Data Fig. 7h). Intriguingly, co-transplanting tumor cells with *Zbtb46*-overexpressed ECs also improved the anti-PD1 treatment outcome (Extended Data Fig. 7i), reinforcing the importance of the EC-specific *Zbtb46* expression in tumor progression. Post-treatment analysis of the TME, lymphoid organs, peripheral blood, and a distal non-tumor organ revealed that the immunostimulatory impact of the systemic nanoparticle treatment was primarily restricted to the TME (Extended Data Fig. 8a,b).

**Fig.4:**
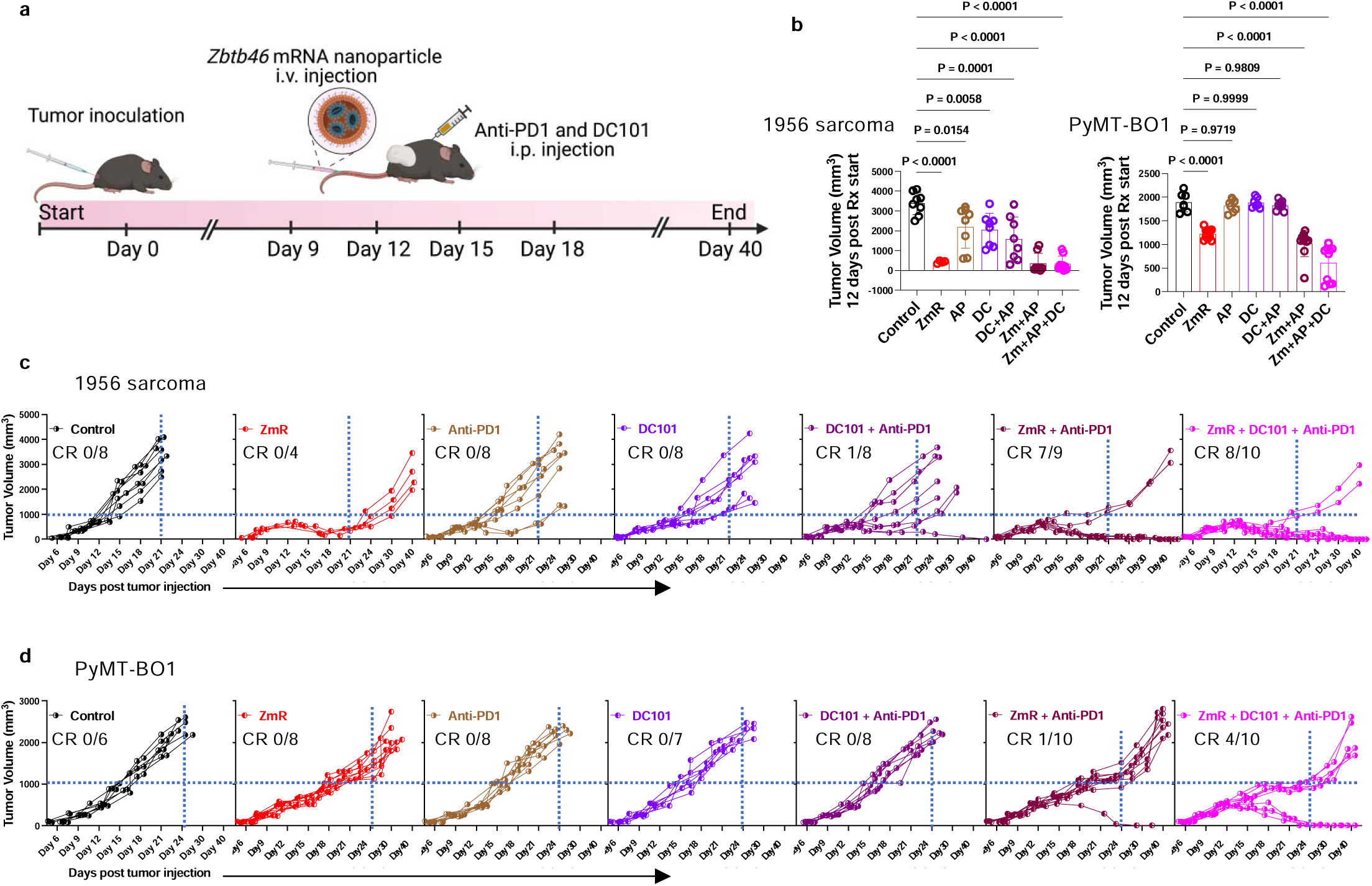
Therapeutic maintenance of *Zbtb46* improves cancer immunotherapy outcome. **a**, Schematics of wild-type tumor-bearing mice treatment with *Zbtb46* mRNA nanoparticle, anti-PD1, and DC101. **b**, Tumor growth of 1956 sarcoma and PyMT-BO1 breast cancer in wild-type mice with *Zbtb46* mRNA nanoparticle (ZmR), anti-PD1 (AP), and DC101 (DC) treatment at 12days post-treatment initiation. n≥6/group. Data are mean±SD. One-way ANOVA with Dunnett’s test. **(c** and **d),** Tumor growth kinetics in (c) 1956 sarcoma and (d) PyMT-BO1 breast cancer in wild-type mice with *Zbtb46* mRNA nanoparticle, anti-PD1, and DC101 treatment. CR=complete remission.

Finally, tumor-derived factors are likely to downregulate *Zbtb46* in the TME. Indeed, tumor-conditioned media (TCM) can downregulate *Zbtb46* expression in vitro (Extended Data Fig. 9a). Among the factors produced by the tumor, we tested a few likely candidates and found that the reactive oxygen species (ROS), prostaglandin E2 (PGE2), retinoic acid (RA), and vascular endothelial growth factors (VEGF) downregulated *Zbtb46* expression in both EC and DC (Extended Data Fig. 9a,b). Pharmacological targeting of the PGE2, RA, and ROS pathways using FDA-approved and commercially available inhibitors modestly prevented tumor-mediated *Zbtb46* suppression both in EC and DC and partially restricted tumor growth (Extended Data Fig. 9c-g), further hinting at the translatability of regulating *Zbtb46* in the tumor.

## Discussion

Tumor vessels and tumor immunity, two crucial TME components, play critical roles in the tumor outcome. Tumor vessels are dilated, tortuous, irregular, and leaky^24, 25^, while tumor immunity is characterized by more immune-suppressive myeloid cell production over dendritic cell generation, known as myeloid lineage skewing^4–7^. Recent studies have revealed an inverse correlation between tumor angiogenesis and tumor immunity^1^. While, clinically, tumor vessel normalization improves tumor immunity and immune checkpoint blockade therapy outcome^10, 11, 26–30^, it is unclear how tumor vessel normalization leads to improved tumor immunity. We provide a common genetic mechanism that can directly control both processes to show how the two seemingly independent processes: tumor angiogenesis and tumor immunity, can be coordinated by *Zbtb46* in cancer.

In mouse tumor models, we found that *Zbtb46* was downregulated in the ECs, DCs, and BM cells. Although *Zbtb46* deficiency did not affect the vasculature or hematopoietic system in non-tumor-bearing mice, tumor-bearing ZKO mice supported more robust tumor growth, accompanied by higher microvascular density, reduced perfusion, augmented vascular leakage and hypoxia, decreased ratios of cytotoxic CTL/Treg and M1/M2 macrophages, diminished cDC1, and increased pro-tumor stroma. Enforcing *Zbtb46* expression in the TME restricted tumor growth accompanied by a remodeling toward an immunostimulatory microenvironment, suggesting a tumor-suppressive role for *Zbtb46* in the TME. Our analysis of publicly available cancer patient datasets also showed that low *ZBTB46* expression is associated with a worse prognosis. Our BM chimera and conditional KO mouse data suggest that both EC- and hematopoietic-specific *Zbtb46* expression are crucial for regulating tumor progression. Our results show that enforcing *Zbtb46* expression leads to less angiogenic and more anti-tumor immune-stimulatory ECs, allowing for increased trans-endothelial leukocyte migration. Although *Zbtb46* deficiency did not disrupt cDC1 antigen processing and presentation, as evidenced by the spontaneous rejection of the 1969 regressive sarcoma tumor model, 1956 progressive sarcoma-bearing ZKO mice had more granulocytic-monocytic progenitors and reduced pre-cDCs in both the BM and spleen. Overexpression of *Zbtb46* in BM cells reversed this trend by generating more dendritic and fewer granulocytic cells, directly impacting tumor growth. It was previously reported that *Zbtb46* targets and represses myeloid lineage-promoting transcription factors, such as *Cebpb*^2, 17^. Our data confirmed that *Cebpb* is a direct target of *Zbtb46*, and *Zbtb46*’s absence resulted in increased transcriptional activity of *Cebpb*. Mechanistically, enforcing *Zbtb46* expression suppresses *Cebpb* transcriptional output and reduces myeloid lineage gene expression in BM cells. We suggest that *Zbtb46* can modulate DC vs. granulocytic lineage output. Our data potentially explains how myeloid lineage skewing is achieved in cancer. Presumably, downregulation of *Zbtb46* and consequent releasing of *Cebpb* repression can lead to more generation of myeloid cells to meet the high demands of myeloid lineage production in emergency conditions, such as cancer^4–7^.

Enforcing *Zbtb46* expression promotes anti-tumor stromal and immune components in the TME, a prerequisite for effective ICB therapy^1, 12^. Indeed, *Zbtb46* mRNA nanoparticles synergized with anti-PD1 treatment to control tumor growth and induce long-term remissions and immunological memory. Not surprisingly, tumor-derived factors suppressed *Zbtb46* expression in vitro, and pharmacological inhibition of these factors partially rescued the tumor-induced *Zbtb46* suppression in vivo. Together, our data uncover a previously unknown tumor-suppressive role of *Zbtb46* in the vascular and hematopoietic system and provide proof-of-concept for using *Zbtb46* maintenance as an effective adjuvant therapy with immunotherapy in cancer management.

## Methods

### TCGA cancer patient dataset analysis for immune microenvironment and survival analysis

We used RNA sequencing data from eighteen non-hematological TCGA tumor types. Cancer types profiled include: Adrenocortical carcinoma (ACC), Bladder Urothelial Carcinoma (BLCA), Breast invasive carcinoma (BRCA), Colon adenocarcinoma (COAD), Cervical squamous cell carcinoma and endocervical adenocarcinoma (CESC), Cholangiocarcinoma (CHOL), Head and Neck squamous cell carcinoma (HNSC), Kidney renal clear cell carcinoma (KIRC), Kidney renal papillary cell carcinoma (KIRP), Liver hepatocellular carcinoma (LIHC), Lung adenocarcinoma (LUAD), Lung squamous cell carcinoma (LUSC), Pancreatic adenocarcinoma (PAAD), Prostate adenocarcinoma (PRAD), Rectum adenocarcinoma (READ), Skin Cutaneous Melanoma (SKCM), and Stomach adenocarcinoma (STAD). RNA sequencing data was downloaded from the GDC pan cancer portal (https://gdc.cancer.gov/about-data/publications/pancanatlas)^42^. Data was processed using the Firehose pipeline with upper quantile normalization. The primary tumor sample was favored for patients with more than one RNA-seq sample. xCell, a gene signatures-based enrichment approach, was used to delineate the enrichment of 64 immune and stromal cell types as described previously^43^. Briefly, the xCell R package was used to generate raw enrichment scores, transform into linear scale, and apply a spillover compensation to derive corrected enrichment scores. The distribution of enrichment scores for patients with high (>50%) and low (<50%) levels of *ZBTB46* expression were compared. *Mann-Whitney* test was used to calculate statistical significance with an alpha value of 0.05.

For survival analysis, the Pathology section of The Human Protein Atlas was accessed through its website (https://www.proteinatlas.org/) and searched for *ZBTB46* gene expression in different tumor types along with the clinical outcome^44, 45^. Briefly, The Cancer Genome Atlas (TCGA) project of Genomic Data Commons (GDC) collects and analyzes multiple human cancer samples. RNA-seq data from 17 cancer types representing 21 cancer subtypes with a corresponding major cancer type in the Human Pathology Atlas were included to allow for comparisons between the protein staining data from the Human Protein Atlas and RNA-seq from TCGA data. The TCGA RNA-seq data was mapped using the Ensembl gene id available from TCGA, and the FPKMs (number Fragments Per Kilobase of exon per Million reads) for each gene were subsequently used for quantification of expression with a detection threshold of 1 FPKM. Based on the FPKM value of each gene, patients were classified into two expression groups, and the correlation between expression level and patient survival was examined. The prognosis of each group of patients was examined by Kaplan-Meier survival estimators, and the survival outcomes of the two groups were compared by log-rank tests.

### Animals

C57BL/6 mice were used as wild-type mice in this study. *Zbtb46^gfp/gfp^* (*Zbtb46* KO) mice were a gift from Kenneth M. Murphy at Washington University in St. Louis. Subsequently, *Zbtb46^gfp/gfp^* mice were crossed with *VEC-Cre, CD11c-Cre, and VAV-Cre* mice (The Jackson Laboratory, Bar Harbor, ME) to generate *VEC-Cre; Zbtb46^gfp/gfp^* conditional KO mice (EC-specific *Zbtb46* KO), *CD11c-Cre; Zbtb46^gfp/gfp^* conditional KO mice (DC-specific *Zbtb46* KO), and *VAV-Cre; Zbtb46^gfp/gfp^* conditional KO mice (Hematopoietic *Zbtb46* KO), respectively. MMTV-PyMT mice were a gift from Mikala Egeblad, Cold Spring Harbor Laboratory, and crossed with *Zbtb46* KO mice to generate *Zbtb46* KO in the presence of MMTV-PyMT transgene (MMTV-PyMT *Zbtb46^gfp/gfp^*). Littermate subjects were used as a control with the different knockout mice. Both male and female mice were used equally in any given experiment except in experiments utilizing both the genetic and orthotopic breast tumor models, where only female mice were used. The ages of the experimental animals were between 10 and 12 weeks.

### Bone marrow (BM) chimeric mice generation

As described previously^46^, wild-type recipient mice (CD45.1) were lethally irradiated with 950rad irradiation. Donor BM from the control (CD45.2) and *Zbtb46* KO (CD45.2) mice was transplanted into the recipient mice retro-orbitally 24-hours post-irradiation. Flow cytometric analysis of the peripheral blood after five months of transplantation confirmed the successful BM reconstitution and generation of ‘hematopoietic *Zbtb46* KO BM chimeric’ mice. Alternatively, lethally irradiated wild-type (CD45.2) and *Zbtb46* KO (CD45.2) recipient mice received BM transplantation from wild-type (CD45.1) donors and generated ‘stromal *Zbtb46* KO BM chimeric’ mice.

### Arterial blood pressure, heart rate, and pressure-diameter measurement in mice

As described previously^47^, *Zbtb46* KO and wild-type littermates were secured under 1.5% isoflurane anesthesia. A Millar pressure catheter (Cat: SPR-671, Millar, Inc.) was introduced to the ascending aorta, and heart rate, arterial systolic, diastolic, and mean blood pressures were recorded using the PowerLab data acquisition system (ADInstruments, Inc.). The ascending aorta and left common carotid artery of the mice were dissected and mounted on metal cannulae in a pressure myograph (Danish Myo Technology). Intravascular pressure was increased from 0 to 175 mmHg in 25 mmHg increments, and the vessel diameter was recorded at each pressure point. The average of 3 measurements at each pressure was reported.

### Mammary tumorigenesis

MMTV-PyMT transgenic mice were utilized to generate a spontaneous model of breast cancer, where MMTV-LTR drives the expression of mouse mammary gland-specific polyomavirus middle T-antigen^23^. Palpable tumors in the mammary gland of the MMTV-PyMT *Zbtb46^gfp/gfp^* (*Zbtb46* KO) mice were measured every week until 21 weeks of age to track the development and progression of tumorigenesis. Tumor volume was calculated by the equation, Volume=(largest diameter) x (smallest diameter)^2^ x 0.5.

### Tumor transplantation studies

1ml of growth factor reduced Matrigel (Cat: 354248; Corning) was mixed with 1ml of cultured LLC-GFP tumor cell suspension (2 x10^6^/ml in PBS); 100μl of the mixture was subcutaneously injected into the back of the mice. 1956- and 1969-sarcoma cells were subcutaneously injected as 1×10^6^ cells in 150μl PBS+Matrigel solution (1:1) per mouse to the flank of the mice. PyMT-BO1 cells were orthotopically injected as 1×10^5^ cells in 50μl PBS+Matrigel solution (1:1) per mouse to the mammary fat pad of the mice, as described previously^48^. Palpable tumors started to develop 4-5 days after transplantation, and tumor growth was measured until the end of the study. For co-transplantation assay, a 1:2 ratio of 1956 sarcoma tumor cells and endothelial cells was mixed in a 1:1 solution of PBS and Matrigel and a total of 2×10^6^ cells in 200μl final volume per mouse was injected into the back of the mice. For overexpression studies, relevant lentiviral particles were intra-tumor injected as 15μl/injection for a viral content of 2×10^6 TU/injection, as many times as indicated in the relevant figures. For in vivo treatment studies, rat IgG2aκ anti-mouse PD1 antibody (Cat: P372, Clone: RMP1-14, Leinco Technologies) was injected intraperitoneally at a dose of 200μg/day, and DC101 (anti-VEGFR2) was injected intraperitoneally at a dose of 40 mg/kg (Cat: BE0060, Bio X cell) (76). Rat IgG2a isotype control at an equivalent dose was used as control.

### Preparation of *Zbtb46* mRNA-p5RHH peptide nanoparticle

For the mRNA nanoparticle treatment study, the *Zbtb46* ORF sequence “ATGAACAACCGAAAGGAAGATATGGAAATCACTTCTCACTACCGGCATCTGCTTCGAGAGCTCAATGAGCAGAGGCAGCACGGAGTCCTCTGTGATGCGTGCGTCGTGGTGGAGGGCAAGGTCTTCAAGGCACATAAGAACGTCTTGCTTGGGAGCAGCCGCTACTTTAAGACGCTCTACTGCCAGGTACAGAAGACATCTGACCAGGCCACCGTCACTCACTTGGACATTGTTACAGCCCAGGGCTTCAAGGCCATTATTGACTTCATGTACTCCGCCCATCTGGCTCTCACTAGTAGGAATGTCATCGAGGTGATGTCAGCTGCCAGCTTCCTACAGATGACTGACATTGTGCAGGCCTGCCATGATTTCATCAAGGCTGCACT GGACATCAGCATAAAGTCAGATGCCTCCGATGAACTCTCAGAATTTGAGATTGGCACCCCAGCCAGCAACAGTACAGAGGCGTTGATCTCAGCTGTGATGGCTGGAAGGAGTATCTCCCCATGGTTGGCTCGGAGAACAAGTCCTGCCAATTCTTCTGGAGACTCTGCCATTGCCAGCTGTCATGAAGGAGGAAGCAGCTATGGGAAGGAGGACCAGGAACCCAAAGCTGATGGCCCTGATGACGTTTCTTCACAGTCTTTGTGGCCTGGAGATGTAGGCTATGGGTCTCTGCGCATCAAGGAAGAACAGATTTCACCATCACATTATGGAGGGAGTGAGCTTCCATCTTCCAAGGACACTGCAATACAGAATTCTTTATCAGAACAGGGTTCTGGGGATGGCTGGCAGCCCACAGGCCGGAGGAAGAATCGGAAAAACAAAGAGACTGTCCGACACATCACCCAGCAGGTGGAGGAGGACAGCCAGGCTGGCTCTCCAGTACCTTCATTCCTACCCACATCGGGATGGCCTTTCAGCAGCCGAGACTCAAATGTAGACCTGACGGTCACTGAGGCCAGCAGCTTGGACAGCCGAGGCGAGAGAGCAGAGCTCTATGCTCACATCGATGAGGGCCTACTAGGAGGAGAAACCAGCTACTTGGGCCCACCCCTCACCCCAGAGAAGGAAGAAGCACTACACCAGGCTACTGCAGTGGCCAATCTTCGTGCTGCACTCATGAGTAAGAACAGTCTGCTGTCACTCAAGGCTGACGTGCTCGGTGATGATGGCTCACTTCTGTTCGAGTACCTGCCCAAAGGTGCCCACTCACTGTCTCGTAAGTGCAAGTTCTGGTGTGTCACTGTGTCTTCCTTTGGTTTAAGCACCTCAGTTCAGCCCTTCAGACCCTGGAGTCACTGA” was made into a modified mRNA transcript with complete substitution of pseudo-U in RNase-free water from TriLink Biotechnologies. CleanCap® EGFP mRNA (Cat: L-7201-100) was used as mRNA control. 8μl (1μg/μl) of the mRNA solution was mixed with 5μl of 20mM p5RHH peptide solution and 187μl of 1x HBSS (Gibco) to prepare the nanoparticle complex and immediately injected into the mouse through the tail vein^21, 29, 37–41^.

### Flow cytometric analysis

Bone marrow (BM) was isolated from the experimental animals by flushing the tibia and femur. Spleen was collected and meshed into a single-cell suspension using the back side of a sterile 5ml syringe plunger. Peripheral blood (PB) was collected by intra-cardiac puncture from the sacrificed animals immediately and processed for making single cells suspension following standard procedure. Lungs were minced and processed by standard enzymatic digestion consisting of Collagenase-I (Cat: LS004194, Worthington). Tumor-draining lymph nodes were isolated, disrupted, and digested with Collagenase-IV (Cat: LS004186, Worthington). Tumor tissues were harvested, minced into fine pieces, and dissociated into single-cell suspensions with an enzymatic digestion buffer consisting of Collagenase-II (for subcutaneous tumors) (Cat: LS004176, Worthington) or Collagenase-III (for breast tumors) (Cat: LS004182, Worthington), along with Dispase-II (Cat: D4693, Millipore Sigma) and Deoxyribonuclease 1 (Cat: LS002139, Worthington). Next, the cell suspensions were incubated with LIVE/DEAD™ Fixable Blue Dead Cell Stain Kit (Cat: L34961) along with different panels of fluorophore-conjugated surface staining antibodies. For subsequent intracellular staining, cell suspensions were fixed and permeabilized using either Foxp3/Transcription Factor Staining Buffer Set (Cat: 00-5523-00, ThermoFisher Scientific) or Intracellular Fixation & Permeabilization Buffer Set (Cat: 88-8824-00, ThermoFisher Scientific) and subsequently stained with intracellular antibodies. Samples were analyzed using either BD LSRFortessa™ X-20 (BD Biosciences) or BD FACSymphony™ A3 (BD Biosciences) and later processed with FlowJo software (BD Life Sciences). CD45-CD31+ ECs and CD45+CD11c+MHCII+ cDCs were FACS sorted using BD FACSAria-II (BD Bioscience). Sorted cells were purity tested by secondary flow cytometry run and later processed for downstream applications total RNA isolation using RNeasy mini kit (Cat:74104, Qiagen) by following the manufacturer’s instructions. Markers used for different cell lineages are given below:

Lineage (Lin): CD3+Ter119+B220+Gr1+

MkP: Lin-Sca1-cKit+CD41+CD150+

GKP: Lin-Sca1-cKit+CD41-CD150-CD16/32+

Pre-cDC: Lin-Sca1-IL7R-MHCII-CD16/32-CD11c+cKit-CD135+

CMP: Lin-Sca1-IL7R-MHCII-CD16/32-CD11c-CD41-CD135+cKit hi

CDP: Lin-Sca1-IL7R-MHCII-CD16/32-CD11c-CD41-CD135+cKit int

cDC: CD45+CD11c+MHCII+

cDC1: CD45+CD11c+MHCII+Xcr1+

CTL: CD45+CD3+CD8+

Treg: CD45+CD3+CD4+CD25+Foxp3+

M1: CD45+CD11b+F4/80+iNOS+

M2: CD45+CD11b+F4/80+CD206+CX3CR1+

B: CD45+B220+

NK: CD45+NK1.1+

EC: CD31+CD45-

### Immunofluorescence studies

Harvested tumors were thinly sliced and fixed in 10% buffered formalin (Cat: 16004-112, VWR), immersed in 30% (w/v) sucrose solution for 48 hours to cryo-protect the tissue, frozen in NEG-50 frozen section medium (Cat: 6502, ThermoFisher Scientific) using liquid nitrogen and 2-methyl butane system, and sectioned in 8μm thickness using a Leica Cryostat microtome (Cat: CM1850, Germany). Afterward, tissue sections were blocked using freshly made blocking buffer (3% essentially IgG-free BSA (Cat: A9085), 0.3% Triton X-100 (Cat: X100), and Fc blocker (Cat: 101301)). Next, sections were incubated with different primary antibodies for 16h at 4°C, followed by visualization with appropriate secondary antibodies. Finally, the sections were counterstained for nuclei with DAPI, cured with ProLong Diamond Antifade mountant, and sealed with nail polish for preservation. To detect intra-tumoral hypoxia in mice, Hypoxyprobe-1 solution (Cat: HP6-100Kit) was intraperitoneally administered 90 minutes prior to sacrifice as 100mg/Kg bodyweight. FITC-conjugated anti-pimonidazole mouse IgG1 monoclonal antibody was used to detect the extent of hypoxia. For the vascular leakage and perfusion experiments, FITC-conjugated 70KD dextran (60mg/Kg bodyweight) (Cat: 46945) and FITC-conjugated lectin (8mg/Kg bodyweight) (Cat: L9381), respectively, were injected through tail vein 15 minutes prior to sacrifice. Hamster anti-mouse CD31 (Clone: 2H8, Cat: MA3105) was used as the primary antibody. AF568 goat anti-hamster (Cat: A-21112) was utilized as the secondary antibody. The sections were examined using the Olympus Fluoview 1200 confocal microscope system and minimally processed with Imaris (Bitplane) software. At least five pictures from every section were processed using ImageJ software (NIH) for quantification purposes.

### Cell lines

Mouse cardiac endothelial cells (MCEC) were purchased from CELLutions Biosystems Inc. (Cat: CLU510). 1956 and 1969 sarcoma cell lines were obtained from Robert D. Schreiber at Washington University in St. Louis. *Zbtb46* overexpressing MCEC and 1956 cell lines were generated by transducing parental MCEC and 1956 cells with *Zbtb46* overexpressing lentiviral particles followed by blasticidin selection (Cat: A1113903, ThermoFisher Scientific). *Zbtb46* knockdown MCEC and 1956 cell lines were generated by transducing the parental MCEC and 1956 cells with lentiviral particles from a set of 5 shRNA clones for *Zbtb46* followed by puromycin selection (Cat: A1113802, ThermoFisher Scientific). GFP expressing Lewis lung carcinoma (LLC cells; Cat: ATCC® CRL-1642™) cells (LLC-GFP) were a gift from Alexander S Krupnick, University of Virginia. PyMT-BO1 cells were obtained from Katherine N Weilbaecher at Washington University in St. Louis. HEK293T cells (Cat: ATCC® CRL-3216™) were purchased from ATCC. All cell lines tested negative for mycoplasma contamination.

### Cell culture

LLC-GFP cells, PyMT-BO1, and HEK293T cells were cultured in DMEM high glucose (Cat: 11965092, ThermoFisher Scientific) growth medium supplemented with 10% (v/v) FBS (Cat: 12103C, Millipore Sigma), 100 unit/ml penicillin-streptomycin (Cat: 15140122, ThermoFisher Scientific). 1956 and 1969 sarcoma cells were cultured in RPMI 1640 growth medium supplemented with 10% (v/v) FBS, 100 unit/ml penicillin-streptomycin, 1% (v/v) L-glutamine (200mM) (Cat: BW17-605E, ThermoFisher Scientific), 1% (v/v) Sodium Pyruvate (100mM) (Cat: BW13-115E, ThermoFisher Scientific), 0.5% (v/v) Sodium Bicarbonate (7.5%w/v stock) (Cat: BW17-613E, ThermoFisher Scientific), and 0.1% (v/v) 2-Mercaptoethanol (Cat: M-6250, Millipore Sigma). All MCEC cell lines were maintained in M199 growth medium (Cat: 11150067, Gibco), supplemented with 20% (v/v) FBS, 100 unit/ml penicillin-streptomycin, and 10mM HEPES (Cat: 15630106, ThermoFisher Scientific). In bone marrow cell-related experiments, bone marrow cells were cultured in Iscove’s Modified Dulbecco’s media supplemented with 10% (v/v) FBS (Cat: 12103C, Millipore Sigma), 100 unit/ml penicillin-streptomycin (Cat: 15140122, ThermoFisher Scientific).

### Lentiviral shRNA and overexpression particle production

pLKpuro lentiviral mouse *Zbtb46* shRNA clones TRCN0000125839 (NM_028125.1-2420s1c1), TRCN0000125840 (NM_028125.1-364s1c1), TRCN0000125841 (NM_028125.1-1358s1c1), TRCN0000125842 (NM_028125.1-321s1c1), and TRCN0000125843 (NM_028125.1-1231s1c1) and mouse Cebpb shRNA clones TRCN0000231407 (NM_009883), TRCN0000231408 (NM_009883), TRCN0000231410 (NM_009883), TRCN0000231411 (NM_009883), and TRCN0000231409 (NM_009883) were purchased from Millipore Sigma. HEK293T cells were transfected with the mentioned shRNA clones or pCSII-EF1-Zbtb46-IRES2-Bsr or pCSII-EF1-Cebpb-IRES2-Bsr constructs along with pCAG-HIVgp and pCMV-VSV-G-RSV-Rev (with a ratio of 4:3:1) by using Calcium Phosphate method. Sixteen hours after transfection, the media was changed, and cells were grown for an additional 48h. Subsequently, the supernatant was harvested and concentrated by Lenti-X-Concentrator (Cat: 631232, Clontech). The virus titer was determined using the Lenti-X™ p24 Rapid Titer Kit (Cat: 632200, Clontech).

### Bone marrow cell differentiation assay

Total bone marrow cells were harvested by flushing out the tibia and femur of the experimental animals. Cells were transfected with different lentiviral particles by spin infection at 800xg for 30min at 32°C in the presence of 2μg/ml polybrene. Cells were later cultured in complete IMDM media with 10ng/ml of Flt3L (Cat: 250-31L, Peprotech), stem cell factor (SCF) (Cat: 250-03, Peprotech), GM-CSF (Cat: 315-03, Peprotech), and G-CSF (Cat: 250-05, Peprotech). Cells were harvested and analyzed by FACS after four days of culture.

### KSL (Lin-cKit+ Sca1+) cell differentiation assay

Total bone marrow cells were harvested by flushing out the tibia and femur of the experimental animals. Cells were then stained with PE/Cy7 conjugated anti-Gr-1 (RB6-8C5), -CD11b (M1/70), -B220 (RA3 6B2), -Ter119 (TER-119) and -CD3 (145-2C11), in combination with APC/Cy7 c-Kit (2B8), PerCp/Cy5.5 Sca1 (E13-161.7), PE CD34 (RAM34) and BV421 CD16/32 antibodies for 40 min on ice. KSL cells (Lin^-^ cKit^+^ Sca1^+^) were sorted on FACS Aria II (BD Biosciences) sorter using 85µm nozzles directly into a round-bottom 96-well plate at a density of 1000 cells/well. Culture media consisted of StemSpan serum-free base medium (StemCell Technologies), 10% FBS, penicillin (50U/mL) and streptomycin (50U/mL), SCF (25ng/ml, PeproTech), FLT3L (20ng/ml, PeproTech), IL3 (1% supernatant), mTPO (20ng/ml, PeproTech), IL6 (10ng/ml, PeproTech), IL11 (10ng/ml, PeproTech), M-CSF (10ng/ml, PeproTech), G-CSF (10ng/ml, PeproTech), and GM-CSF (10ng/ml, PeproTech). Cells were harvested and analyzed by FACS after six days of culture.

### Bone marrow monocyte enrichment

Tibia and femur of the experimental animals were harvested and flushed, and later monocytes were enriched from the total bone marrow cells using the Monocyte Isolation Kit (BM) from Miltenyi Biotec (Cat: 130-100-629) through a negative selection process following the manufacturer’s instruction. This enriched population contained mostly monocytes and precursor lineage-negative cells.

### Chromatin Immunoprecipitation (ChIP)-qPCR Assay

As described previously^49^, enriched monocyte and lineage-negative cells from bone marrow were cross-linked with 1% formaldehyde and lysed with cell membrane lysis buffer, followed by further incubation with nuclear membrane lysis buffer. For the *Zbtb46* overexpression system, before proceeding to lysate preparation, after the isolation of the enriched population, *Zbtb46*-HA overexpression lentiviral particles were used to transduce the cells by spin infection at 800xg for 30min at 32°C in the presence of 2μg/ml polybrene and then incubated at 37°C for 24hours in complete IMDM media. The lysate was incubated with either an anti-CEBPb antibody (Cat: 23431-1-AP, Proteintech) or anti-HA antibody (Cat: ab9110, Abcam) followed by protein A/G sepharose beads (Cat: sc-2002, Santa Cruz Biotechnology). The beads were washed to isolate immunoprecipitated DNA fragments that were later subjected to qPCR. For anti-ZBTB46 experiment Cebpb peak-1 enrichment (DNA location chr20:50189053-50189756; corresponding mouse region: chr2:167687660-167688195)^50^ was assessed by qPCR using primer-set forward sequence “CCCCAGCTCAGCAGATAACA” and reverse sequence “AGGCTTCTCAGGTGATTGCG”. For anti-CEBPb experiment, Cebpb peak-1 enrichment (DNA location chr2:167687679-167688023)^51^ was assessed by qPCR with primer-set forward sequence “AAGGGCACAGGGAGATGTCA” and reverse sequence “GGTGTTGCTCAACCTTCGGT”; Csf3r peak-6 enrichment (DNA location chr4:126017669-126017995)^51^ was assessed by qPCR with primer-set forward sequence “GACAACGCTGGCACTTTTGTA” and reverse sequence “TGTGCAAGCAGGTCATTGTG”.

### Intra-tumoral transfer of enriched bone marrow monocytes

Enriched monocytes and precursor lineage-negative cells from CD45.1 wild-type donor mice were transduced with either empty vector-mCherry or Zbtb46-mCherry lentiviral particles by spin infection at 800xg for 30min at 32°C in the presence of 2μg/ml polybrene. Later, as described previously^52^, the cells were resuspended in PBS as 1×10^5^ cells/50μl and injected intra-tumor into the day-10 post-transplant 1956 sarcoma tumor-bearing CD45.2 wild-type recipient mice. Tumor volumes were measured periodically and harvested at day-15 post-transplantation for flow cytometric analysis to track donor-derived (CD45.1+) transduced (mCherry+) cell polarization into either macrophage or dendritic cells.

### Tube formation assay

Overnight serum-starved endothelial cells were plated on Matrigel (Cat: 96992, Corning) coated 24-well plates (3×10^4^ cells/well) and incubated at 37°C for 6 hours before taking pictures with a Leica DFC 310 FX microscope system as described before^21^. The angiogenesis analyzer module of ImageJ software (NIH) was used to quantify the total tube length, number of loops, and number of branches.

### T cells trans-endothelial migration assay

As described before^29^, T cell migration potential across an endothelial barrier in vitro was assessed using a QCM™ Leukocyte Trans-endothelial Migration Colorimetric Assay kit (Cat: ECM557, Millipore Sigma) following the manufacturer’s instruction. T cells used in the study were isolated from the spleens harvested from 1956 sarcoma subcutaneous tumor-bearing wild-type mice with the APC CD45 (Cat: 103112, Clone: 30-F11), PE CD3 (Cat:100206, Clone: 17A2), BV650 CD4 (Cat: 100469, Clone: GK1.5), PerCP/Cy5.5 CD8a (Cat: 100734, Clone: 53-6.7). Cytotoxic T cells were sorted using BD FACSAria™ II (BD Biosciences) as CD45^+^CD3^+^CD4^-^CD8^+^ population. Endothelial monolayers were treated with either TNFα (20ng/ml) for 12 hours and washed with PBS before placing the harvested T cells on the endothelial monolayer. The relative abundance of migrated T cells was calculated by measuring the absorption of the samples at 450nm following the WST-1 reagent staining.

### Quantitative real-time reverse transcription PCR (qRT-PCR)

cDNA was prepared with qScript™ cDNA SuperMix (Cat: 101414-106, VWR) according to the manufacturer’s protocol. Gene expression was measured by quantitative real-time RT-PCR using primers detailed in Supplementary Table 1.

### Bulk RNA sequencing

RNA sequencing data from WT and *Zbtb46* overexpressing MCECs was pre-processed and analyzed for differentially expressed genes using edgeR^53^. Genes with fewer than five counts in at least three samples were filtered out. Count data was normalized to account for varying library sizes. Normalized data was fit to a negative binomial generalized log-linear model, and differentially expressed genes were extracted. In-built functions for gene ontology analysis were used to identify GO terms enriched in WT and *Zbtb46* overexpression MCECs.

### Statistical Analysis

GraphPad Prism 8 software was used for performing statistical analysis and generating graphs/plots. Data are presented as mean with standard deviation (SD) for all the measurements. Statistical significance was determined by two-tailed unpaired Student’s t test (for two groups) and one-way analysis of variance (ANOVA) with Dunnett’s or Tukey’s multiple comparison test, as appropriate (for more than two groups). Non-parametric tests were used for non–log-transformed gene expression data from the TCGA database. *P < 0.05* was considered statistically significant.

### Graphical presentations

Graphical schematics presented in this work were prepared with BioRender.com.

### Data availability

Sequencing data is available as GSE226087. Source data are provided with this paper.

### Study approval

Animal husbandry, generation, handling, and experimentation were performed in accordance with protocols approved by the Institutional Animal Care and Use Committee of Washington University School of Medicine in St. Louis.

## Acknowledgements

We thank K. M. Murphy at Washington University in St. Louis for the *Zbtb46^gfp/gfp^* (*Zbtb46* KO) mice and M. Egeblad at Cold Spring Harbor Laboratory for MMTV-PyMT mice. We thank our colleagues at Washington University, R.D. Schreiber for 1956 and 1969 sarcoma cells, K. Lavine for MCEC cells and K. Weilbaecher for PyMT-BO1-GFP-Luc cells. We also want to thank A. S. Krupnick at the University of Virginia for providing LLC-GFP cells. We thank Washington University Center for Cellular Imaging (WUCCI) and Pathology FACS core for providing access to the light microscopes and FACS facility, respectively. We also thank GTAC@MGI for performing the RNA sequencing.

## Author contributions

A.U.K. and K.C. conceived the idea, designed the experiments, and wrote the manuscript. A.U.K. performed all experiments and analyzed data. M.S. analyzed bulk RNA sequencing data and TCGA patient dataset. M.K. helped with the ChIP-qPCR experiments. J.W. and K.K. helped with bone marrow chimeric mice generation and analysis. C.M.H. helped with the cardiovascular measures of the ZKO mice. H.P. and S.A.W. provided peptides for the nanoparticles. X.W. and D.H.F. helped with the *Zbtb46*-lentiviral constructs. K.C. provided overall supervision and coordinated all the experimental activities. All authors approved the final manuscript.

