## Supplementary Figures for "*Zbtb46* coordinates angiogenesis and immunity to control tumor outcome"

Campus Box 8118, 660 S. Euclid Ave. St. Louis, MO, 63110-1093, USA

Key words: Cancer, Tumor microenvironment, Dendritic cell, Endothelial cell, *Zbtb46*, *Cebpb*, Myeloid lineage skewing

**Extended data figures and tables**

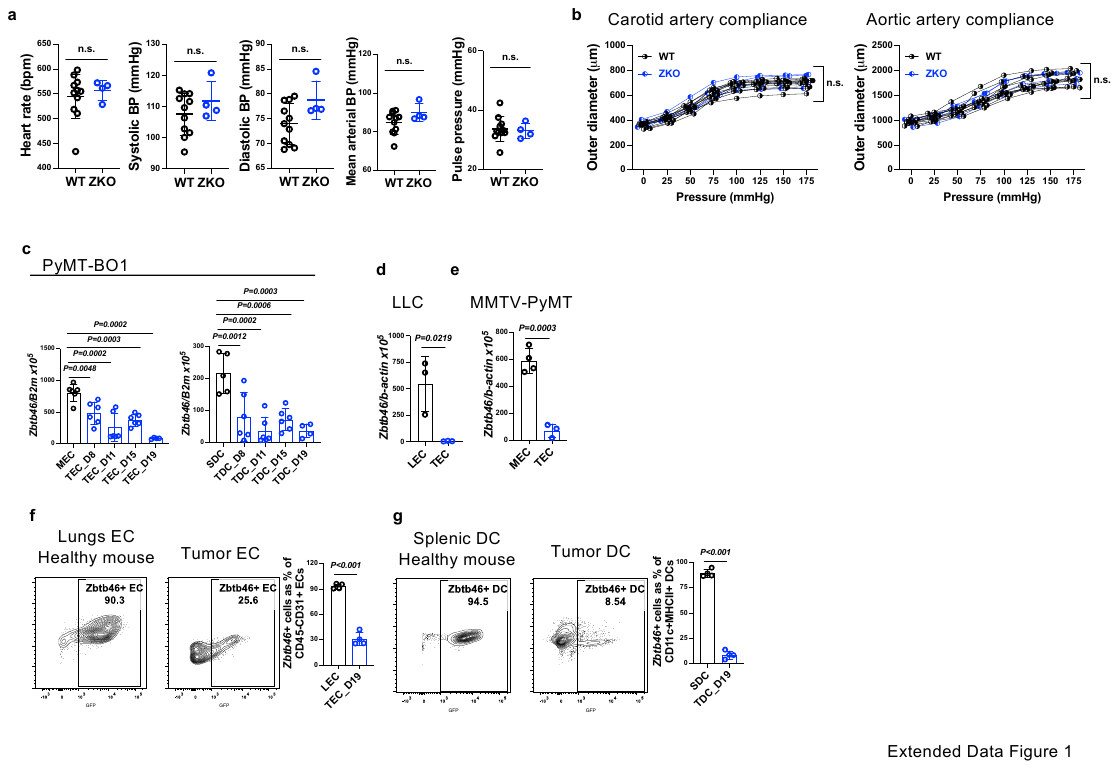

**Extended Data Fig.1: *Zbtb46* expression is downregulated in tumor**

**a-b**, Analysis of (a) vital cardiovascular parameters and (b) pressure-diameter measurements in the *Zbtb46* KO and wild-type mice. n≥4/group. Data are mean±SD. Student’s t-test (a) and One-way ANOVA with Tukey’s multiple comparison test (b). **c**, *Zbtb46* mRNA expression in (left) CD31+CD45- ECs and (right) CD45+CD11c+MHCII+ DCs isolated from the mammary gland (MEC) and Spleen (SDC) of the healthy wild-type mice and from the tumors (TEC and TDC) of the PyMT-BO1-bearing wild-type mice at indicated days post-transplantation. n≥3/group. Data are mean±SD. One-way ANOVA with Dunnett’s test. **d-e**, *Zbtb46* mRNA expression in CD31+CD45- ECs isolated from the lungs (LEC) and mammary gland (MEC) of the healthy wild-type mice and from the tumors (TEC) of the (b) LLC carcinoma and (c) MMTV-PyMT bearing mice. n≥3/group. Data are mean±SD. Student’s t-test. **f-g**, Analysis of ZBTB46+(GFP+) (f) CD31+CD45- ECs and (g) CD45+CD11c+MHCII+ DCs isolated from the lungs and spleen of the healthy and tumors of the 1956-bearing *Zbtb46^gfp/+^* mice. n≥3/group. Data are mean±SD. Student’s t-test.

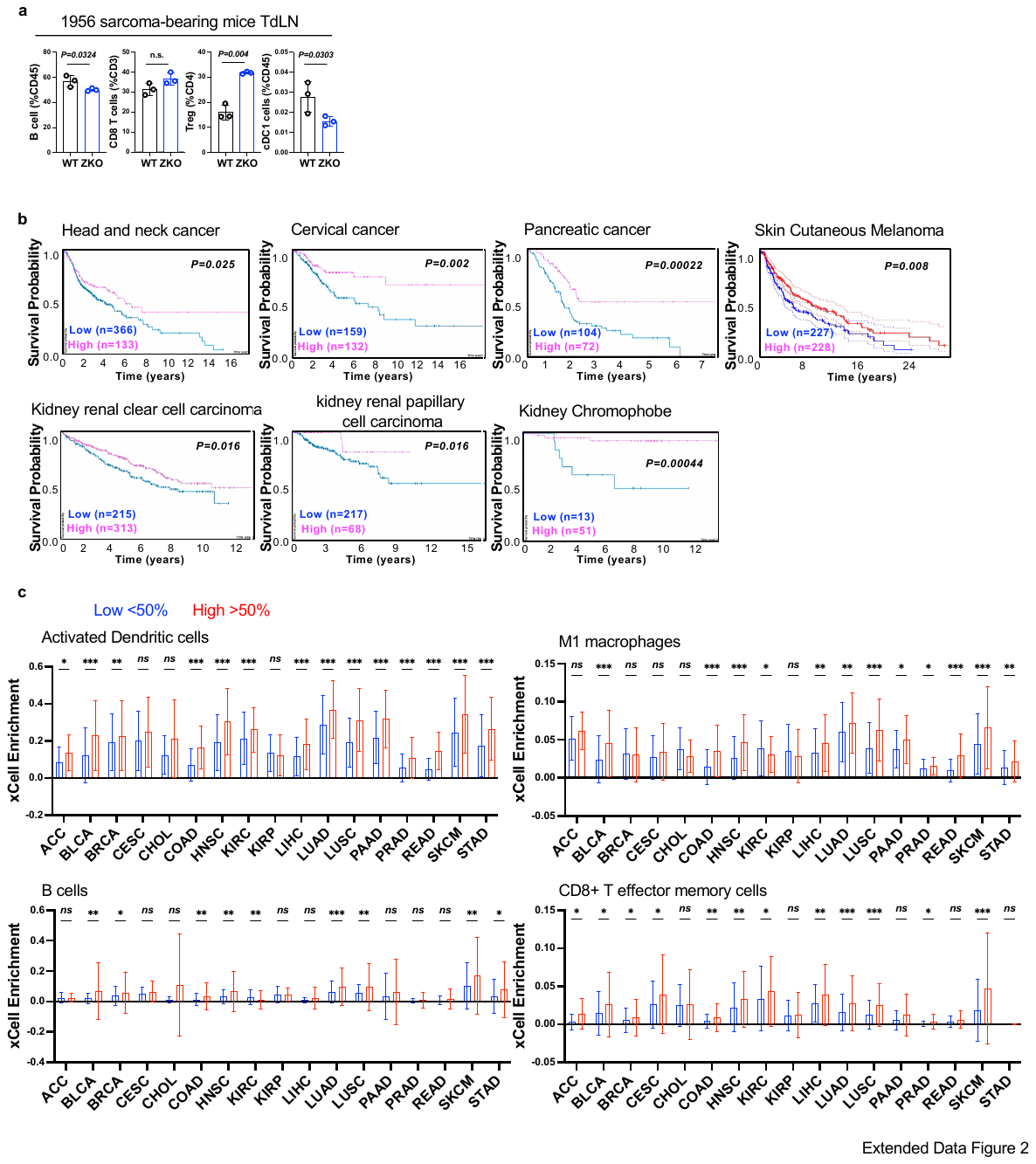

**Extended Data Fig.2: *Zbtb46* is a tumor suppressor**

**a**, Measurements of B, CD8T, Treg, and cDC1 cells in the tumor-draining lymph nodes of 1956 sarcoma-bearing mice. n=3/group. Data are mean±SD. Student’s t-test. **b**, Overall survival of patients with indicated cancer separated by *ZBTB46* expression as high (>50th percentile) and low (<50th percentile). Survival data were derived from publicly available clinical records of TCGA patients. Log ranked test was used for survival analysis. **c**, Presence of activated dendritic cells, M1 macrophages, B cells, and CD8+ T effector memory cells in high vs. low *ZBTB46* expressing tumors in patients from the TCGA database analyzed with the CIBERSORT algorithm. Data are mean±SEM. Student’s t-test. *p<0.05, **p<0.01, ***p<0.001, ns=not significant.

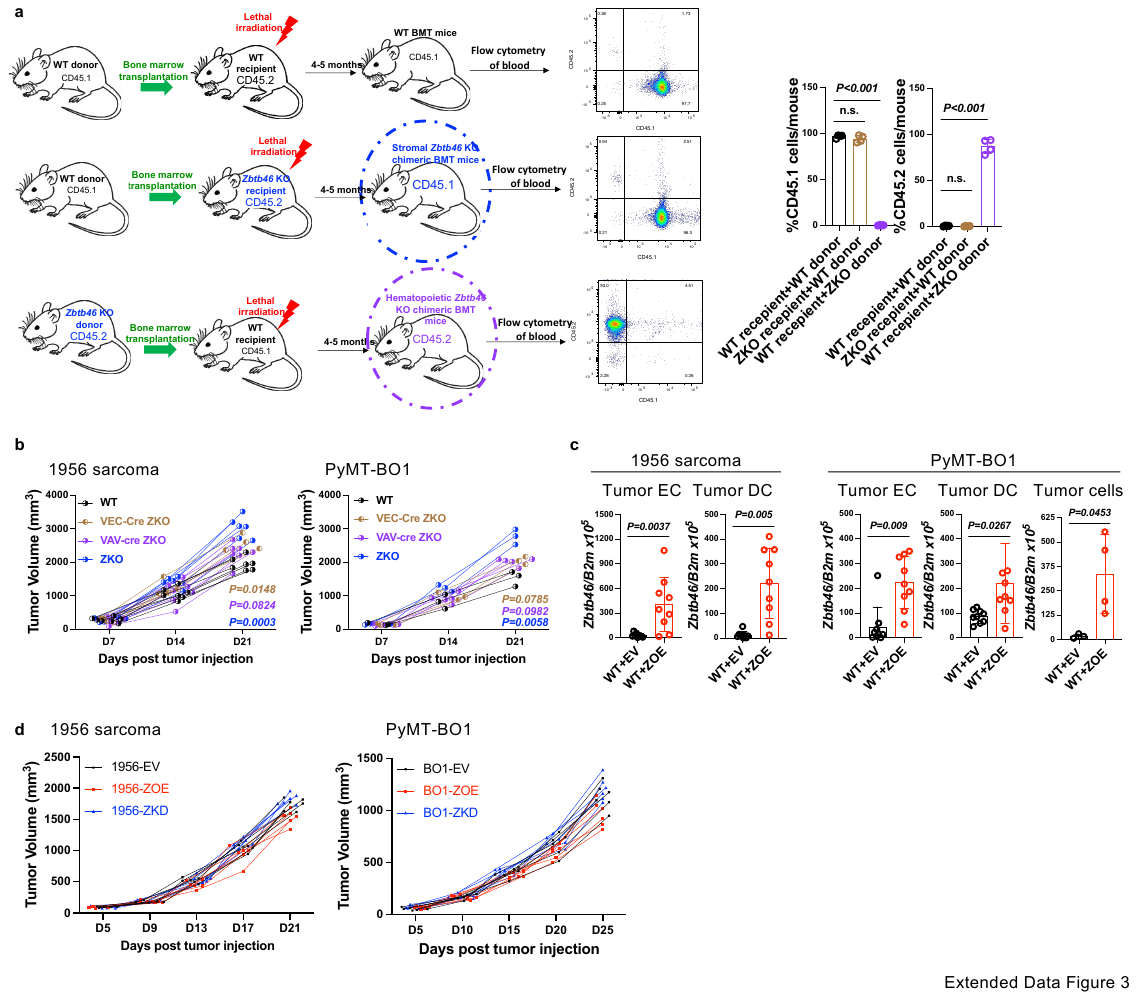

**Extended Data Fig.3: Both endothelial and hematopoietic *Zbtb46* contribute to the suppression of tumor progression**

**a**, Schematic of stromal and hematopoietic *Zbtb46* KO bone-marrow chimera mice generation and flow cytometric analysis of CD45.1 (for stromal KO) or CD45.2 (for hematopoietic KO) repopulation in the peripheral blood. n≥3/group. Data are mean±SD. One-way ANOVA with Dunnett’s test. **b**, Tumor growth of 1956 sarcoma and PyMT-BO1 breast cancer in wild-type, VEC-cre *Zbtb46* KO, VAV-cre *Zbtb46* KO, and *Zbtb46* KO mice. n≥3/group. One-way ANOVA with Dunnett’s test at endpoint compared to wild-type mice. **c**, *Zbtb46* mRNA expression in CD31+CD45- ECs, CD45+CD11c+MHCII+ DCs, and GFP+ tumor cells (TC) isolated from the tumors of the 1956 sarcoma and PyMT-BO1-bearing wild-type mice with empty vector (EV) or *Zbtb46* (ZOE) lentiviral overexpression construct intra-tumor treatment. n≥3/group. Data are mean±SD. Student’s t-test. **d**, Tumor growth of 1956 sarcoma and PyMT-BO1 with empty vector (1956-EV and BO1-EV) or *Zbtb46* (1956-ZOE and BO1-ZOE) lentiviral overexpression or *Zbtb46* shRNA construct expression (1956-ZKD and BO1-ZKD) in wild-type mice. n≥4/group. One-way ANOVA with Dunnett’s test at endpoint.

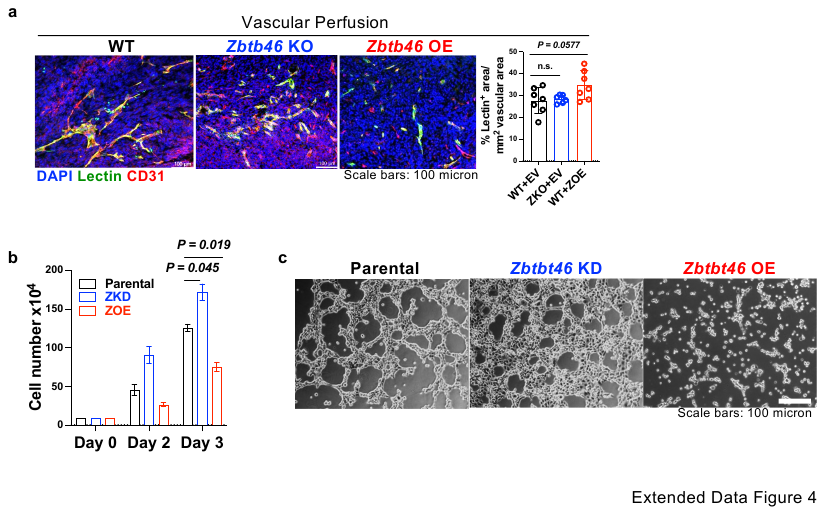

**Extended Data Fig.4: *Zbtb46* maintains endothelial cells quiescent in the tumor context**

**a**, Representative images and quantification for vascular perfusion as measured by the FITC-lectin binding to vessels in the 1956 sarcoma tumor tissue of wild-type mice with either empty vector (WT+EV) or *Zbtb46* (WT+ZOE) or of *Zbtb46* KO mice with empty vector (ZKO+EV) lentiviral overexpression construct (intra-tumor) treatment. n≥4/group. Data are mean±SD. One-way ANOVA with Dunnett’s test. **b**, Analysis of proliferation by counting cell numbers of the cultured parental, *Zbtb46* knockdown (ZKD), and *Zbtb46* overexpressing (ZOE) MCEC cells. n≥3/group. Data are mean±SD. One-way ANOVA with Dunnett’s test. **c**, Representative images and quantifications from the Matrigel tube formation assay with parental, *Zbtb46* knockdown (ZKD), and *Zbtb46* overexpressing (ZOE) MCEC cells.

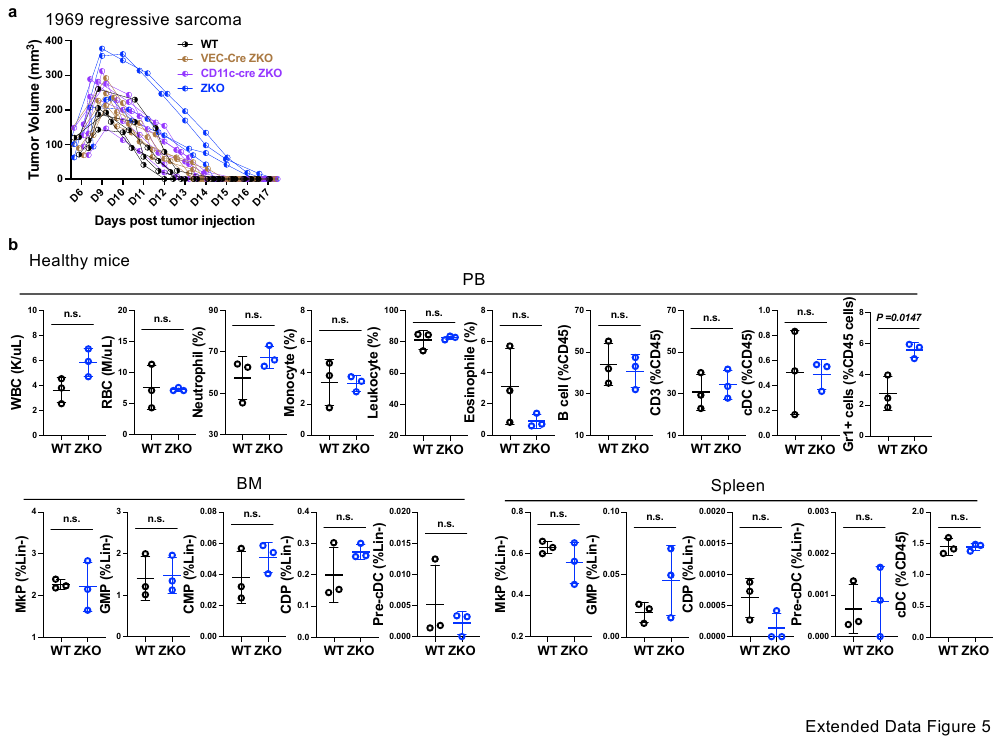

**Extended Data Fig.5: *Zbtb46* KO hematopoietic system is similar to wild-type in homeostasis but more immunosuppressive in the tumor context**

**a**, Tumor growth of 1969 regressive sarcoma in wild-type, VEC-cre *Zbtb46* KO, CD11c-cre *Zbtb46* KO, and *Zbtb46* KO mice. n≥4/group. **b**, Hemavet analysis and flow-cytometric analysis of peripheral blood (PB), bone marrow (BM), and spleen in the wild-type and *Zbtb46* KO mice in healthy condition. Markers for different cell lineages are provided in the Methods section. n≥3/group. Data are mean±SD. Student’s t-test.

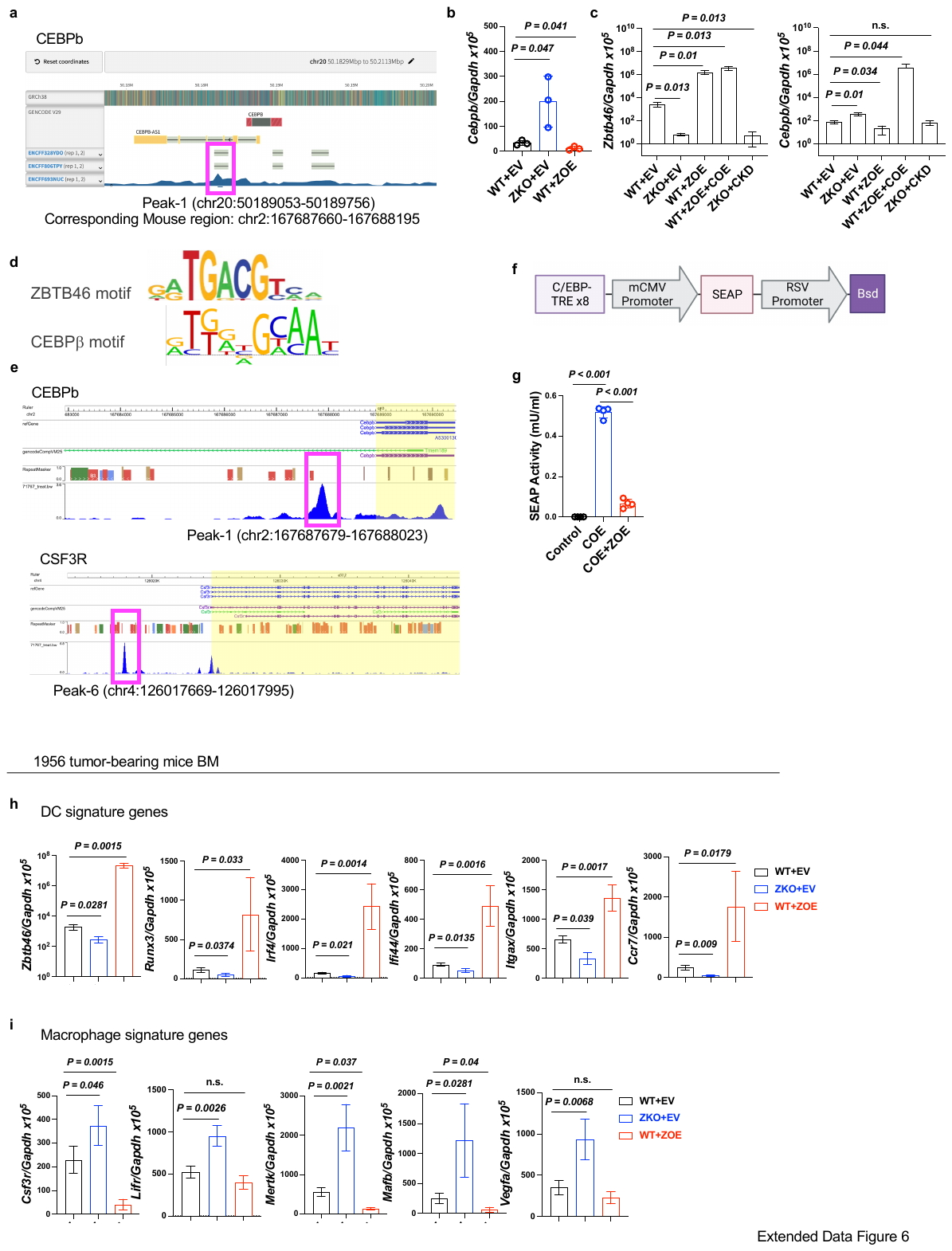

**Extended Data Fig.6: *Zbtb46* sustains dendritic cell lineage generation while suppressing myeloid lineages in the tumor context**

**a**, Genomic snapshots depicting the ZBTB46 binding regions at the indicated genomic loci. **b**, *Cebpb* mRNA expression in the bone marrow cells of wild-type or *Zbtb46* KO mice with either empty vector (EV) or *Zbtb46* (ZOE) lentiviral overexpression. n=3/group. Data are mean±SD. One-way ANOVA with Dunnett’s test. **c**, *Zbtb46* and *Cebpb* mRNA expression in the bone marrow cells of wild-type or *Zbtb46* KO mice with either empty vector (EV), or *Zbtb46* (ZOE), or *Cebpb* shRNA constructs (CKD), or *Cebpb* (COE) lentiviral overexpression. n=3/group. Data are mean±SD. One-way ANOVA with Dunnett’s test. **d**, Consensus motif for ZBTB46 and CEBPB ChIP sequences. **e**, Genomic snapshots depicting the CEBPB binding regions at the indicated genomic loci. **f**, Schematics of reporter lentivector core expression cassette for CEBP signaling pathway. **g**, Chemiluminescence measurement of SEAP activity in the reporter-only (control), or reporter with *Cebpb* overexpressed (COE), or reporter with *Cebpb* and *Zbtb46* overexpressed (COE+ZOE) assay system. Data are mean±SD. One-way ANOVA with Dunnett’s test. **h-i**, Analysis of a few (h) DC and (i) macrophage signature genes in 1956 sarcoma-bearing bone marrow cells from wild-type and *Zbtb46* KO mice with empty vector (EV) or *Zbtb46* (ZOE) overexpression. n=3/group. Data are mean±SD. One-way ANOVA with Dunnett’s test.

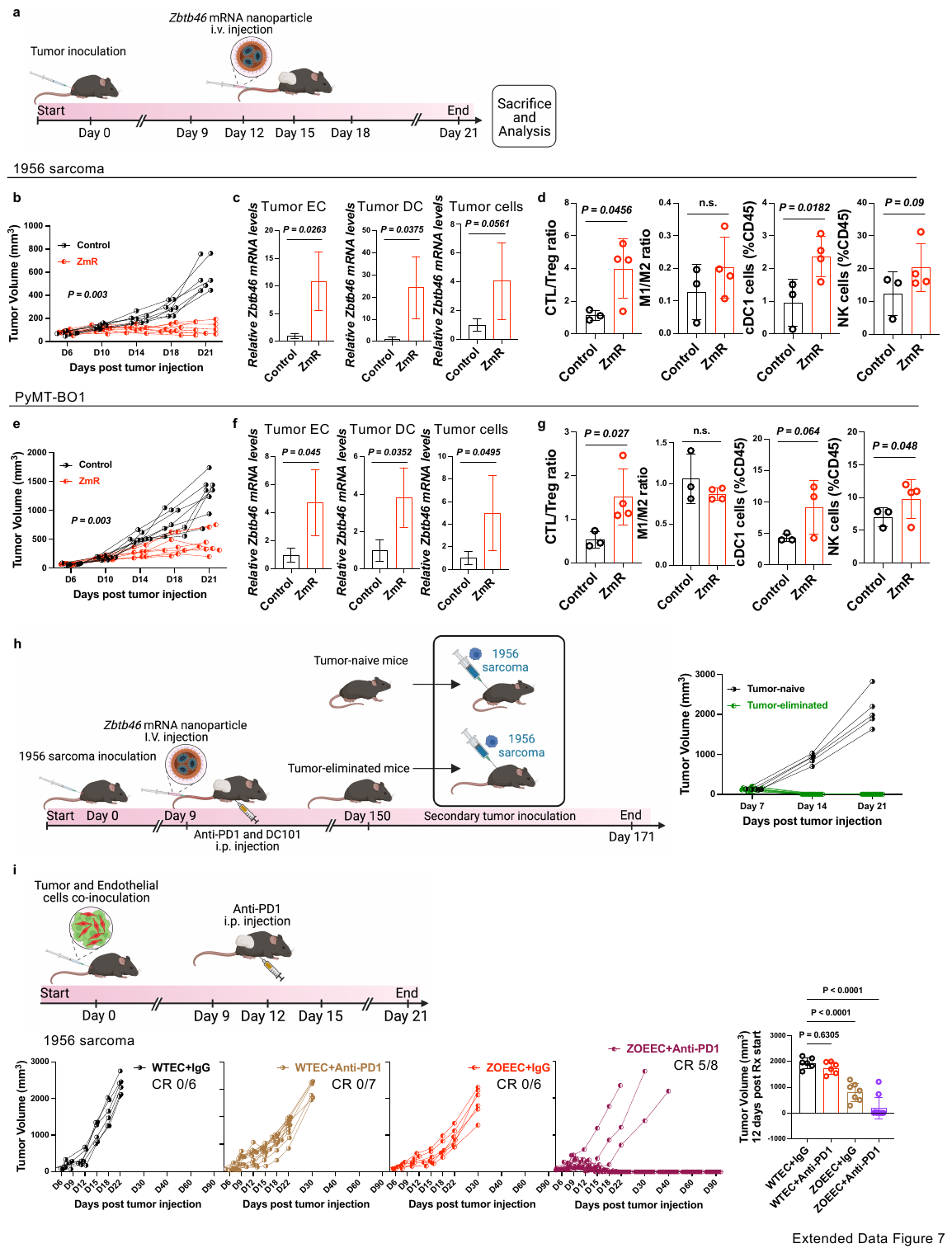

**Extended Data Fig.7: Systemic *Zbtb46* mRNA nanoparticle treatment restricts tumor growth and enhances outcomes of anti-PD1 immunotherapy**

**a**, Schematics of wild-type tumor-bearing mice treatment with *Zbtb46* mRNA nanoparticle. **b-g**, Tumor growth kinetic (b, e), *Zbtb46* mRNA expression (c, f), and immune microenvironment (d, g) of 1956 sarcoma (b-d) and PyMT-BO1 breast cancer (f-g) in wild-type mice with *Zbtb46* mRNA nanoparticle (ZmR) treatment. n≥3/group. Data are mean±SD. Student’s t-test at endpoint. **h,** Schematic and tumor growth in the treatment-responded 1956 sarcoma tumor-eliminated mice challenged with secondary 1956 sarcoma transplantation. **i,** Schematics and tumor growth of 1956 sarcoma cells co-transplanted with parental endothelial cells (WTEC) or *Zbtb46* overexpressed endothelial cells (ZOEEC) with IgG or anti-PD1 treatment. n≥5/group. CR= Complete Remission. One-way ANOVA with Dunnett’s test at 12days post-treatment initiation.

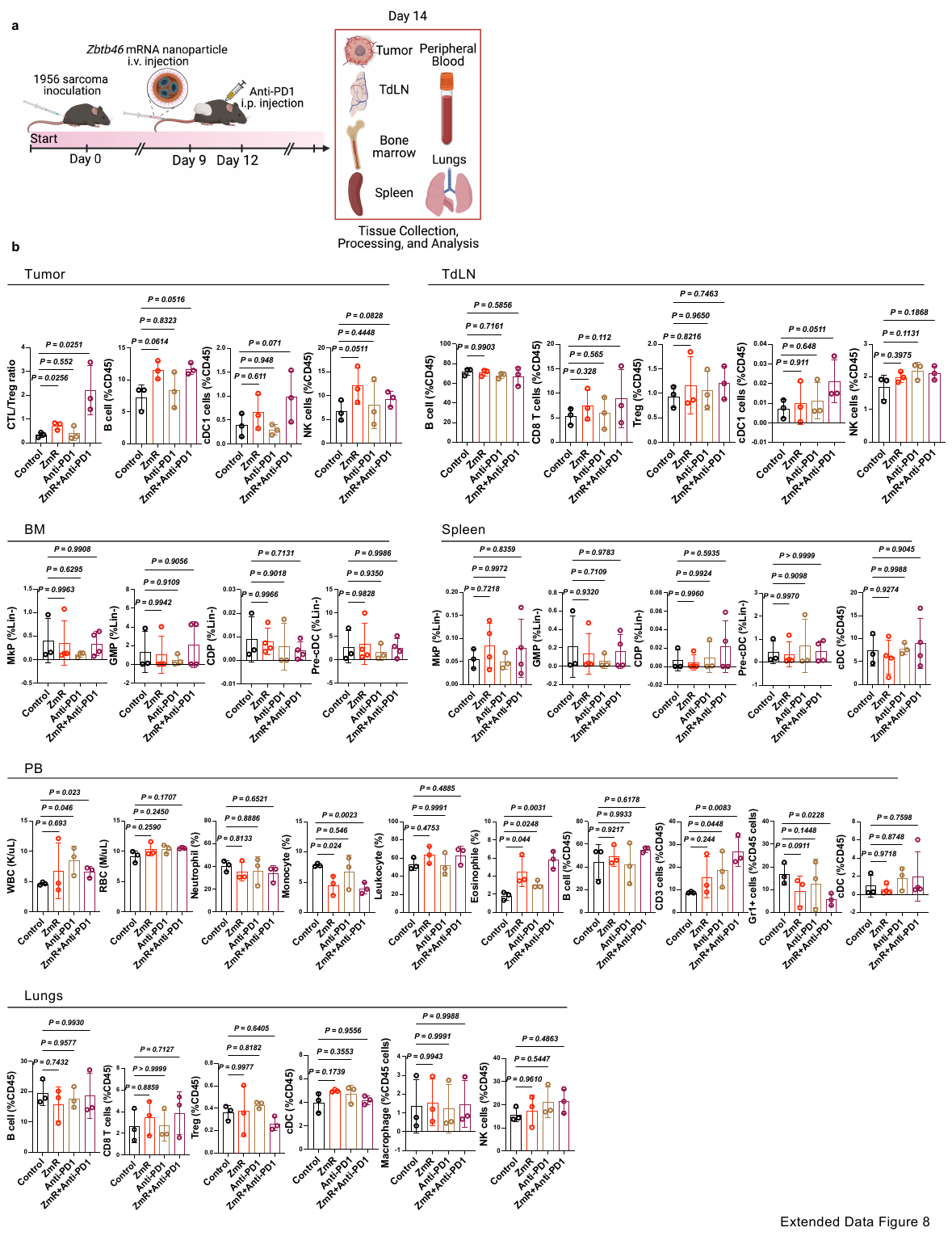

**Extended Data Fig.8: Systemic *Zbtb46* mRNA nanoparticle and anti-PD1 treatment leads to immunostimulatory TME**

**a**, Schematics of analysis of TME and different distal organs from 1956 sarcoma-bearing wild-type mice with *Zbtb46* mRNA nanoparticle (ZmR) and anti-PD1 treatment. **b**, Hemavet and flow cytometric analysis of tumor, tumor draining lymph node (TdLN), bone marrow (BM), spleen, peripheral blood (PB), and lungs for immune components and precursor cells. Markers for the cell lineages are provided in the methods section. n≥3/group. One-way ANOVA with Dunnett’s test.

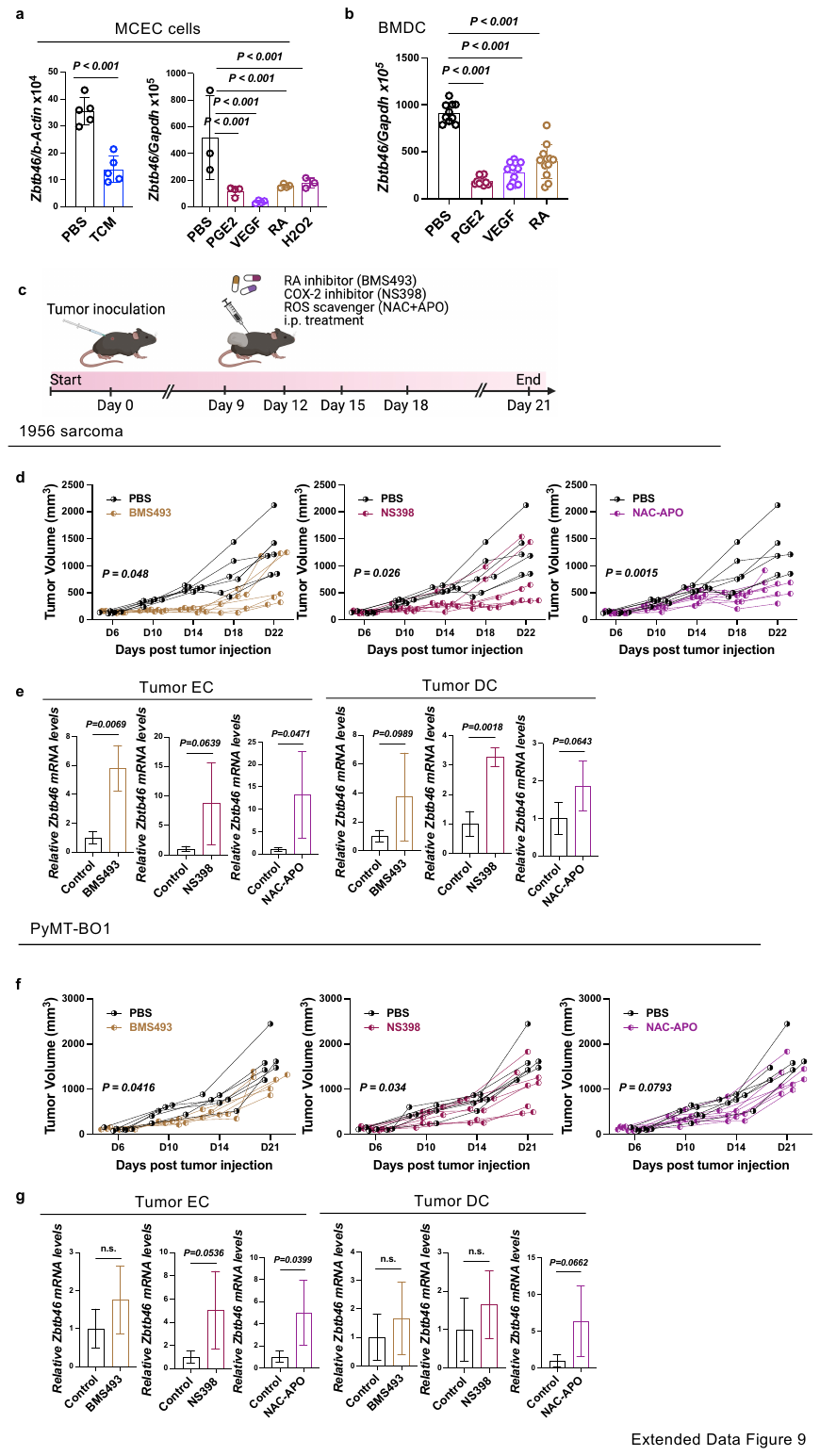

**Extended Data Fig.9: Tumor-derived factors suppress *Zbtb46***

**a-b**, *Zbtb46* mRNA expression in (a) MCEC cells and (b) bone marrow-derived dendritic cells (BMDC) with indicated treatments for 24hours. n≥3/group. One-way ANOVA with Dunnett’s test. **c**, Schematics depicting treatment plan for tumor-bearing mice. **d-g**, Tumor growth kinetic (d, f) and *Zbtb46* mRNA expression (e, g) of 1956 sarcoma (d-e) and PyMT-BO1 breast cancer (f-g) in wild-type mice with BMS493 (inverse pan-retinoic acid receptor agonist), NS398 (selective cyclooxygenase-2 inhibitor), and NAC+APO (ROS scavengers) treatment. n≥6/group. Data are mean±SD. Student’s t-test at endpoint.

Supplementary table 1: Sequences of Primers used in the study

| Gene | Forward Sequence | Reverse Sequence | Description |
| --- | --- | --- | --- |
| *Cebpb* peak-1 | CCCCAGCTCAGCAGATAACA | AGGCTTCTCAGGTGATTGCG | Anti-ZBTB46 ChIP-qPCR |
| *Cebpb* peak-1 | AAGGGCACAGGGAGATGTCA | GGTGTTGCTCAACCTTCGGT | Anti-CEBPB ChIP-qPCR |
| *Csf3r* peak-6 | GACAACGCTGGCACTTTTGTA | TGTGCAAGCAGGTCATTGTG | Anti-CEBPB ChIP-qPCR |
| *Zbtb46* | ATCACTTCTCACTACCGGCAT | AAGACGTTCTTATGTGCCTTGAA | qRT-PCR |
| *Cebpb* | CGCCTTATAAACCTCCCGCT | TGGCCACTTCCATGGGTCTA | qRT-PCR |
| *Cd5l* | GATCGTGTTTTTCAGAGTCTCCA | TGCAGTCAACCCCTTGAATAA G | qRT-PCR |
| *Runx3* | CAGGTTCAACGACCTTCGATT | GTGGTAGGTAGCCACTTGGG | qRT-PCR |
| *Irf4* | TCCGACAGTGGTTGATCGAC | CCTCACGATTGTAGTCCTGCTT | qRT-PCR |
| *Ifi44* | AACTGACTGCTCGCAATAATGT | GTAACACAGCAATGCCTCTTGT | qRT-PCR |
| *Itgax* | CTGGATAGCCTTTCTTCTGCTG | GCACACTGTGTCCGAACTCA | qRT-PCR |
| *Ccr7* | TGTACGAGTCGGTGTGCTTC | GGTAGGTATCCGTCATGGTCTTG | qRT-PCR |
| *Lifr* | TACGTCGGCAGACTCGATATT | TGGGCGTATCTCTCTCTCCTT | qRT-PCR |
| *Mertk* | CAGGGCCTTTACCAGGGAGA | TGTGTGCTGGATGTGATCTTC | qRT-PCR |
| *Mafb* | TTCGACCTTCTCAAGTTCGACG | TCGAGATGGGTCTTCGGTTCA | qRT-PCR |
| *Vegfa* | CTGCCGTCCGATTGAGACC | CCCCTCCTTGTACCACTGTC | qRT-PCR |
| *Csf3r* | CTGATCTTCTTGCTACTCCCCA | GGTGTAGTTCAAGTGAGGCAG | qRT-PCR |
